# When Does a Pathway Score Preserve a Mediated Effect? Aggregation Geometry, Minimax Adequacy, and Gap Inference for High-Dimensional Mediation

**DOI:** 10.64898/2026.10.02.756279

**Authors:** Subir Hait

## Abstract

High-dimensional mediators are routinely collapsed into low-dimensional scores before mediation analysis, but aggregation can change the mediated estimand rather than merely reduce noise. We characterize when a linear score preserves the full mediated effect. Exact preservation for every outcome model occurs precisely when the score contains the covariance-adjusted exposure-predictive mediator direction, the classical sufficient reduction for exposure under a Gaussian inverse-regression model. We introduce a distribution-free minimax adequacy loss equal to the fraction of exposure-relevant mediator information discarded by the score and exactly linked to overlap recovered by aggregation. A split-sample inverse-Wishart correction yields unbiased estimators of retained and full exposure information in moderate dimensions. Under spherically symmetric outcome-effect benchmarks, relative aggregation error has a Cauchy location-scale profile, and paired influence-function inference targets the realized gap for an observed outcome. In simulations, paired interval coverage ranged from 93.3% to 97.5%. In an outcome-free TCGA lung adenocarcinoma audit of 1,314 smoking-responsive genes, the pooled gene-mean loss was 0.811 (descriptive 95% gene-bootstrap interval 0.797–0.823), versus a dimension-only random-subspace benchmark of 0.913; retaining promoter and non-promoter means separately reduced pooled loss to 0.625. Independent half-sample screening was unstable, but top-100 and top-250 cross-screen pooled gene-mean losses ranged from 0.827 to 0.932. The framework separates structural adequacy, benchmark fragility, and outcome-specific discrepancy.

## 1 Introduction

Mediation analyses with omics mediators face a dimensionality problem that is usually solved by aggregation. Methylation at hundreds of CpGs is averaged within a promoter or gene; transcripts are summarized into pathway activity scores; imaging voxels are reduced to principal components. Pathway software reports direction-weighted activation scores (Krämer et al., 2014), and singular-value summaries of gene sets are a standard pathway-level statistic (Tomfohr et al., 2005). Epigenetic clocks are themselves fixed linear combinations of CpGs (Horvath, 2013; Lu et al., 2019), and they are widely used as mediators. Methods that estimate indirect effects through many mediators simultaneously have developed in parallel (Zhang et al., 2016; Huang and Pan, 2016; Chén et al., 2018; Derkach et al., 2019; Luo et al., 2020).

The implicit justification for aggregation is statistical: fewer mediators, less variance. This paper asks a prior, causal question. *When does the indirect effect through an aggregate equal the indirect effect through the mediators it summarizes?* If it does not, a pathway-level analysis answers a different question from the one its conclusions are phrased in, and it may do so with misleadingly narrow uncertainty.

The question is related to, but distinct from, three literatures. Work on principal directions of mediation (Chén et al., 2018) and on transformed mediators (Huang and Pan, 2016) *constructs* lowdimensional directions that carry the indirect effect; we instead *audit* directions chosen by someone else, typically without reference to the outcome. Work on ambiguous manipulations (Spirtes and Scheines, 2004), causal coarsening (Chalupka et al., 2016), and causal abstraction (Rubenstein et al., 2017; Beckers and Halpern, 2019) asks when a macro-variable supports well-defined causal claims; we quantify how the mediated estimand changes when it does not, in a setting where coarsening is routine. Finally, work on constructed measures (VanderWeele, 2022) emphasizes that a composite and its components need not share causal roles; we give that concern an exact form for mediation. The present paper also extends an earlier analysis of aggregation distortion in multilevel mediation (Hait, 2026) to the high-dimensional setting.

Our contributions are as follows. Section 2 gives an exact projection identity, links exact preservation to sufficient dimension reduction under a Gaussian inverse model, introduces a distribution-free minimax adequacy loss, and derives a finite-sample unbiased split-sample estimator of the retained exposure information when the relevant covariance matrices are estimable. Section 3 generalizes the Cauchy adequacy profile from Gaussian to spherically symmetric outcome-effect benchmarks and turns the profile into probabilities of practically meaningful errors. Section 4 develops paired influence-function inference for the realized gap *D*_*B*_, including a sample-splitting protocol for scores learned from the data. Sections 5 and 6 evaluate the outcome-free diagnostics and gap inference, respectively. Section 7 first performs a genome-wide, outcome-free audit of annotation-based gene summaries and then illustrates outcome-specific gaps using smoking, gene methylation, and gene expression. The aim is not to advocate a universally “best” summary but to make the estimand change caused by a proposed summary visible before, and testable after, an outcome is introduced.

The analysis is related to dimension reduction but has a different target. High-dimensional mediation methods use transformed or sparse mediator directions to facilitate testing and estimation (Huang and Pan, 2016; Chén et al., 2018; Zhao et al., 2020); sufficient-dimension-reduction methods seek low-dimensional summaries retaining regression information (Cook, 2007; Cook and Forzani, 2008; Luo et al., 2017). Under the normal inverse-regression model used by principal fitted components, a direction proportional to Σ^*−*1^*α* is the one-dimensional sufficient reduction of the mediator vector for predicting the exposure. That classical fact is not claimed as new here. What is new is the mediation consequence: an externally chosen score preserves the full mediated effect for *every* outcome model if and only if its span contains that sufficient exposure-reduction direction, and the amount omitted is exactly connected to worst-case mediation distortion and recovered overlap. We therefore use the term *exposure-predictive mediator direction* for Σ^*−*1^*α*. The phrase “mediator propensity score” is avoided because it is already used in mediation weighting methods for a different object.

## 2 Setup and aggregation geometry

### 2.1 Model and estimands

Let *A* be an exposure, *W* ∈ ℝ^*q*^ pre-exposure covariates, *M* ∈ ℝ^*p*^ mediators, and *Y* an outcome. We work with the linear structural model

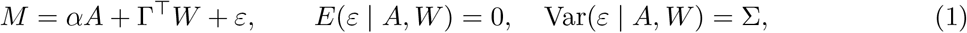

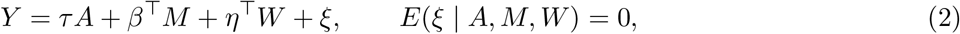

with Σ positive definite and *v*_*A*_ = Var{*A* − *E*^*\**^(*A* | *W*)} *>* 0, where *E*^*\**^ denotes linear projection. Under the usual no-unmeasured-confounding conditions for the exposure–mediator, exposure– outcome, and mediator–outcome relations given *W*, the indirect effect of a unit change in *A* through *M* (natural or interventional, which coincide here; VanderWeele et al., 2014; Vansteelandt and Daniel, 2017) is IE_*M*_ = *α*^*⊤*^*β*, the direct effect is DE_*M*_ = *τ*, and the total effect is TE = *τ* + *α*^*⊤*^*β*.

An *aggregation* is a fixed *p* × *k* matrix *B* of full column rank, *k* ≤ *p*, with aggregate *Z* = *B*^*⊤*^*M* . Gene means, promoter means, sign-weighted scores, and principal-component scores are all of this form once their weights are fixed.

#### Definition 1

(Aggregated effects). Let DE_*Z*_ be the coefficient of *A* in the linear projection of *Y* on (*A, Z, W*), and define IE_*Z*_ = TE − DE_*Z*_ and the aggregation gap *D*_*B*_ = IE_*M*_ − IE_*Z*_.

IE_*Z*_ is exactly what an analyst estimates by running a product-of-coefficients or difference-method mediation analysis with *Z* in place of *M* . When (*ε, ξ*) is jointly Gaussian, projections coincide with conditional expectations and IE_*Z*_ is the interventional indirect effect through *Z*; in the general case of (1)–(2) it is the corresponding projection estimand.

### 2.2 The projection identity

#### Proposition 1

(Projection identity and reallocation). *Under* (1)*–*(2),

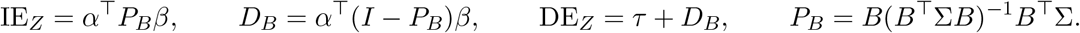

*In particular* IE_*Z*_ + DE_*Z*_ = IE_*M*_ + DE_*M*_ = TE *for every B*.

*P*_*B*_ is idempotent with range col(*B*); it is the oblique projection onto col(*B*) along ker(*B*^*⊤*^Σ). The last statement is the practically important one: aggregation does not discard mediated effect, it relabels it as direct. An aggregate analysis reporting “no mediation through the pathway” and a sizable direct effect can be fully consistent with strong mediation through the pathway’s components.

Writing *a* = Σ^*−*1*/*2^*α, b* = Σ^1*/*2^*β*, and Π_*B*_ = Σ^1*/*2^*B*(*B*^*⊤*^Σ*B*)^*−*1^*B*^*⊤*^Σ^1*/*2^, the orthogonal projection onto col(Σ^1*/*2^*B*), the gap is *D*_*B*_ = ⟨(*I* −Π_*B*_)*a*, (*I* −Π_*B*_)*b*⟩ and hence |*D*_*B*_| ≤ ∥(*I* −Π_*B*_)*a*∥ ∥(*I* −Π_*B*_)*b*∥. Sign reversal, IE_*M*_ IE_*Z*_ *<* 0, is possible even with *k* = 1: with Σ = *I, B* = *e*_1_, *α* = (1, 1)^*⊤*^ and *β* = (1, −2)^*⊤*^, one has IE_*M*_ = −1 but IE_*Z*_ = 1.

### 2.3 Exact preservation

#### Proposition 2

(Exact preservation). *(i) D*_*B*_ = 0 *for every β if and only if* Σ^*−*1^*α* ∈ col(*B*). *(ii) D*_*B*_ = 0 *for every α if and only if β* ∈ col(*B*). *(iii) For given* (*α, β*), *D*_*B*_ = 0 *if and only if* (*I* − Π_*B*_)*a* ⊥ (*I* − Π_*B*_)*b*.

#### Corollary 1

*The one-dimensional scores α*^*⊤*^Σ^*−*1^*M and β*^*⊤*^*M are each exactly adequate*.

#### Remark 1

The condition *α* ∈ col(*B*), which may seem natural, is *not* sufficient for exact preservation. With *B* = *α* = *e*_1_ one obtains IE_*Z*_ = (Σ*β*)_1_*/*Σ_11_, which differs from IE_*M*_ = *β*_1_ whenever Σ is not diagonal. Section 3 shows that *α* ∈ col(*B*) removes *systematic* distortion only.

### 2.4 Connection to sufficient dimension reduction

The direction in Proposition 2 has a classical inverse-regression interpretation. Principal fitted components and related sufficient-dimension-reduction methods obtain forward-regression information by modelling the distribution of predictors given the response (Cook, 2007; Cook and Forzani, 2008). In the present notation, the “response” for that inverse regression is the exposure and the “predictors” are the mediators.

#### Proposition 3

(Sufficient exposure reduction). *Assume in addition that*

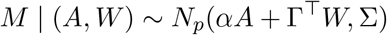

*with constant positive-definite* Σ. *Conditional on W, the likelihood of A given M depends on M only through*

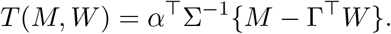

*Thus T is a one-dimensional sufficient reduction of M for predicting A given W; under the usual non-degeneracy condition for the exposure, the corresponding central subspace is* span(Σ^*−*1^*α*). *Consequently, a fixed aggregate B*^*⊤*^*M preserves the mediated effect for every outcome direction β if and only if its column space contains this sufficient exposure-reduction direction*.

Proposition 3 clarifies the relation to existing dimension-reduction theory. The sufficient direction itself is not a new object: it is the one-index specialization of the normal inverse-regression reductions developed by Cook and Forzani. The new statement is the causal-decomposition consequence. Containing the sufficient exposure reduction is exactly the universal preservation condition for mediation, while failing to contain it creates the minimax loss and overlap trade-off developed below. In this sense, sufficient dimension reduction supplies the direction and the present analysis supplies its mediation interpretation.

### 2.5 Adequacy and overlap

#### Proposition 4

(Adequacy–overlap trade-off). *Let κ* = *α*^*⊤*^Σ^*−*1^*α and κ*_*B*_ = *α*^*⊤*^*B*(*B*^*⊤*^Σ*B*)^*−*1^*B*^*⊤*^*α. Then*

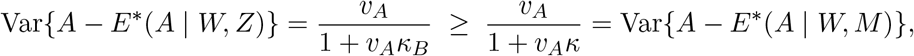

*with equality if and only if* Σ^*−*1^*α* ∈ col(*B*). *Moreover the coefficient vector of M in E*^*\**^(*A* | *W, M*) *is proportional to* Σ^*−*1^*α*.

Proposition 4 has two consequences. First, the exactly adequate one-dimensional score of Corollary 1 is, up to scale, the linear predictor of the exposure from the mediators given covariates; we call Σ^*−*1^*α* the *exposure-predictive mediator direction*. Second, any aggregate that preserves the mediated effect for every outcome model leaves exactly as little residual exposure variation as the full mediator vector, so it inherits the full vector’s positivity problems. When mediators nearly determine exposure, as in our data (Section 7.1), no aggregate can be simultaneously adequate and substantially improve overlap.

### 2.6 A distribution-free minimax adequacy loss

Exact preservation is binary, while most scientific scores are inexact. A useful outcome-free measure should therefore quantify how much of the exposure-relevant mediator direction is excluded without assigning any distribution to *β*.

#### Definition 2

(Minimax adequacy loss). For *κ >* 0, define

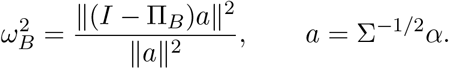

If *α* = 0, set *ω*_*B*_ = 0 by convention.

#### Proposition 5

(Minimax and overlap representations). *Let κ* = *α*^*⊤*^Σ^*−*1^*α and*

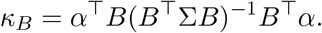

*For κ >* 0,

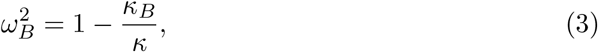

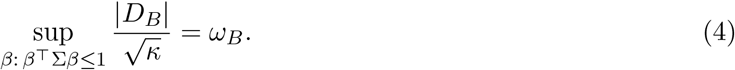

*Thus* 0 ≤ *ω*_*B*_ ≤ 1, *and ω*_*B*_ = 0 *if and only if B is exactly adequate for every outcome model. If*

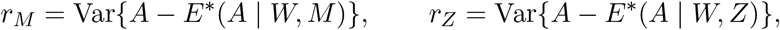

*then*

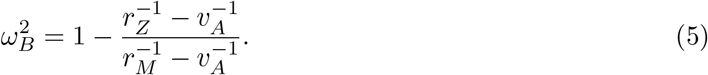

Equation (4) gives *ω*_*B*_ a direct causal interpretation: among outcome directions with one unit of residual-mediator scale, it is the largest possible aggregation gap relative to the exposure signal available in the full mediator vector. Equation (3) gives the same number an exposure-prediction interpretation. The fraction 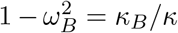 is the share of exposure-relevant mediator information retained by *B*. Consequently, adequacy and overlap do not merely move in opposite directions qualitatively; they are two views of the same geometry.

#### A random-subspace benchmark

The magnitude of 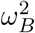 should be interpreted relative to the dimension of the retained subspace. In whitened coordinates, suppose the *k*-dimensional aggregation subspace is drawn uniformly at random and independently of the fixed exposure direction *a*.

Rotational symmetry then gives

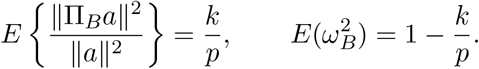

We use 1 − *k/p* as a geometric reference for how much information a dimension-*k* reduction would retain absent alignment with the exposure direction. It is exact for a uniformly random subspace in whitened space; it is not a null distribution for raw random weights in the original correlated mediator coordinates. A scientifically constructed score is informative to the extent that it improves on this dimension-only benchmark.

This result also explains why an exposure-blind summary can look statistically convenient. A small *κ*_*B*_ leaves substantial exposure variation after adjustment for *Z*, but the improvement in overlap is purchased by a large worst-case adequacy loss. Conversely, driving *ω*_*B*_ toward zero forces *κ*_*B*_ toward *κ* and hence recovers the same overlap difficulty as the full mediator set. The relation is invariant to invertible reparameterizations of the columns of *B* and therefore applies to a pathway subspace rather than to a particular choice of basis.

### 2.7 Estimating retained exposure information

The identity 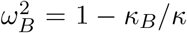is useful only if the two information quantities can be estimated. A direct plug-in estimator is upward biased because it uses an estimated exposure-effect vector on both sides of an inverse sample covariance. In moderate dimensions, a simple two-sample construction removes both sources of first-order finite-sample bias exactly under Gaussian mediator errors.

Randomly split the observations into two groups (stratified by exposure when appropriate). In group *h* ∈ {1, 2}, regress *M* on (1, *A, W*) and let 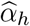 be the coefficient vector of *A, R*_*h*_ the residual matrix, and *d*_*h*_ the residual degrees of freedom. Define *d* = *d*_1_ + *d*_2_ and

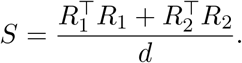

For a fixed rank-*k* aggregation *B*, define

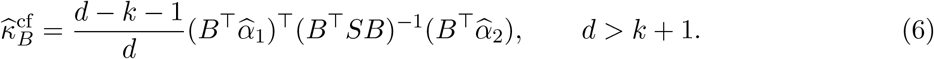

When *d > p* + 1, the full information is estimated by

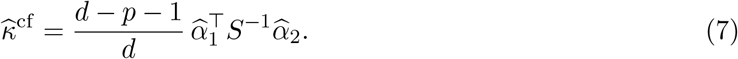

#### Proposition 6

(Unbiased information estimators). *Under the Gaussian mediator model of Proposition 3, with a fixed full-rank design in each split and fixed B, the estimators in* (6) *and* (7) *satisfy*

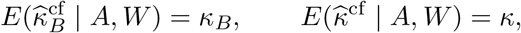

*whenever the stated degrees-of-freedom conditions hold*.

The corresponding estimator

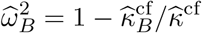

is a ratio and is therefore not exactly unbiased, but it is consistent when both information estimators are. Finite-sample values can fall slightly outside [0, 1], especially when *κ* is weak; we recommend reporting the untruncated estimate, together with 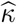, rather than hiding this instability by clipping. Repeating the stratified split and averaging 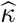 and 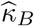 before forming the ratio reduces split sensitivity. For uncertainty, a stratified nonparametric bootstrap should repeat the complete splitting and score-construction procedure. The exact correction requires *p < d* − 1 for the full *κ*; this is feasible for gene-level audits with tens of CpGs even when the genome-wide mediator vector itself has *p* ≫ *n*.

### 2.8 Invariance and monotonicity

The adequacy problem concerns a subspace, not the particular columns used to represent that subspace. This is useful both conceptually and computationally.

#### Proposition 7

(Basis invariance and nested subspaces). *Let B have full column rank*.

1. *If R is any invertible k* × *k matrix and B*^*\**^ = *BR, then* 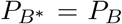. *Consequently D*_*B*_, *ω*_*B*_, *m*_*B*_, *γ*_*B*_, *and the aggregated indirect effect are invariant to a change of basis within the same aggregation subspace*.
2. *If* col(Σ^1*/*2^*B*_1_) ⊆ col(Σ^1*/*2^*B*_2_), *then* 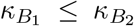 *and* 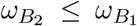 . *Thus enlarging a fixed aggregation subspace cannot worsen the distribution-free minimax loss*.

The second statement is intentionally limited to *ω*_*B*_. The realized *D*_*B*_ need not move monotonically toward zero as directions are added because positive and negative omitted contributions can cancel. Likewise the Cauchy location *m*_*B*_ need not be monotone for arbitrary oblique projections. A larger score vector therefore improves the worst-case geometric guarantee while not necessarily improving a particular outcome estimate at every intermediate dimension. This distinction is visible in the constructed scenarios in the Supplement: adding PCs usually helps, but cancellation can make a smaller summary accidentally closer for one chosen *β*.

Basis invariance also clarifies what is meant by a multi-score mediator. If a pathway is represented by three PCs, rotating those three PCs by an orthogonal matrix does not change the mediation estimand because the column space is unchanged. More generally any nonsingular rescaling or mixing of the three scores leaves *P*_*B*_ unchanged. The scientific decision is therefore the retained subspace, not the naming or normalization of its coordinates.

### 2.9 An exposure–mediator interaction extension

The projection geometry extends to a simple exposure–mediator interaction, but only under explicit additional conditions. Let *A* ∈ {0, 1} and retain the mediator model

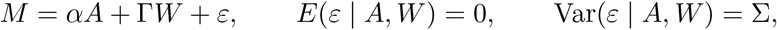

where the residual covariance Σ is the same in both exposure arms. Replace (2) by

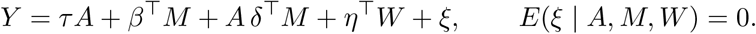

For *a* ∈ {0, 1}, define the exposure-level indirect contrast

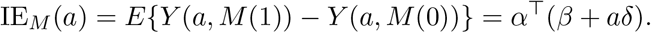

For the aggregated analysis, the conditional mean must include *Z* = *B*^*⊤*^*M*, the interaction *A* × *Z*, and, whenever Γ ≠ 0, the corresponding *A* × *W* terms. The latter are required because *Aδ*^*⊤*^*E*(*M* | *A, Z, W*) contains an *A* × *W* component after conditioning on *Z*. Under this correctly specified interacted model, define IE_*Z*_(*a*) analogously from the fitted aggregate model. The same projection calculation then gives

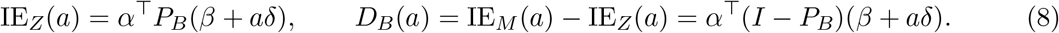

Thus the aggregation gap can change with the exposure level, with *D*_*B*_(1) − *D*_*B*_(0) = *α*^*⊤*^(*I* − *P*_*B*_)*δ*. Universal exact preservation at both exposure levels still holds if and only if Σ^*−*1^*α* ∈ col(*B*). If the residual covariance differs by exposure arm, a single *P*_*B*_ no longer describes both arms and the identity above must be replaced by an arm-specific projection analysis. The homogeneous model is the special case *δ* = 0.

### 2.10 What exact adequacy requires for common scores

Proposition 2 becomes especially informative when translated into the summaries analysts actually use. Let *c* = Σ^*−*1^*α* denote the exposure-predictive mediator direction. Exact adequacy for every outcome model requires *c* ∈ col(*B*). The following cases show why seemingly reasonable summaries can fail for different reasons.

#### Equal-weight mean

For a single standardized mean, *B* ∝ **1**_*p*_. Exact preservation requires

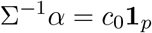

for some scalar *c*_0_. Equivalently, after accounting for mediator covariance, every component must contribute the same coefficient to the linear prediction of the exposure. Equal exposure coefficients *α*_*j*_ are not enough unless the covariance has the appropriate symmetry. This explains why a gene mean can be inadequate even when all CpGs move in the same direction with smoking: correlation changes which linear combination contains the exposure information relevant to the mediation decomposition.

#### Grouped means

Suppose mediators are divided into *G* disjoint groups and *B* contains their indicator vectors, possibly normalized by group size. Exact preservation requires *c* to be piecewise constant across those groups. Separate hyper- and hypomethylation means therefore improve on a single signed score when the exposure-predictive coefficients are approximately homogeneous within the two classes, but their adequacy is a statement about Σ^*−*1^*α*, not simply about the signs of *α*. The same logic applies to promoter and non-promoter means, cell-type region summaries, and pathway submodules.

#### Exposure-weighted score

A tempting construction is *B* = *α*. Because *α* ∈ col(*B*), Corollary 2(i) gives *m*_*B*_ = 0: under the spherical benchmark there is no systematic relative bias. Yet exact adequacy requires Σ^*−*1^*α* ∝ *α*, which holds if and only if *α* is an eigenvector of Σ. Unless the exposure direction aligns with a residual-covariance eigenvector, correlation rotates the exactly adequate direction away from the raw exposure coefficients. The exposure-weighted score can therefore have zero average distortion and substantial unpredictable distortion at the same time. This is precisely what happens in the semi-synthetic TCGA population.

#### Sign score

For a one-dimensional sign score with weights *b*_*j*_ ∈ {+*c*_+_, −*c*_*−*_}, exact preservation requires Σ^*−*1^*α* to be proportional to that two-level vector. The score discards not only magnitudes of the exposure coefficients but also covariance-induced reweighting. Consequently a sign score may be interpretable and transportable yet geometrically fragile. Its adequacy should be reported rather than inferred from the fact that its signs agree with the exposure associations.

#### Principal components

If *B* contains eigenvectors of the residual covariance Σ, exact preservation is equivalent to *α* lying in their span. To see this, write Σ = *V* Λ;*V* ^*⊤*^ and *α* = *V d*. Then

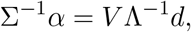

so the zero pattern across eigen-directions is the same for *α* and Σ^*−*1^*α*. Retaining a set *J* of residual-covariance PCs gives

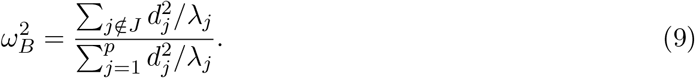

Thus the minimax diagnostic weights exposure signal in low-variance directions more heavily, whereas Corollary 2(iii) gives

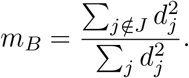

The two diagnostics answer different questions: *ω*_*B*_ uses the residual-covariance metric that governs exact preservation, while *m*_*B*_ uses the outcome-effect benchmark on the standardized mediator scale. A PC may explain substantial mediator variance while retaining little of the exposure-predictive direction, and conversely a low-variance PC can matter greatly for exact adequacy.

Empirical PCs introduce another distinction. Corollary 2(iii) and Equation (9) use eigenvectors of the residual covariance after removing (*A, W*). PCs formed from the raw mediator matrix incorporate between-exposure and covariate variation and need not be eigenvectors of Σ. This is why the empirical PC profiles in the application need not obey 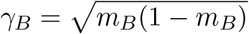 exactly.

#### Published biomarkers and clocks

For a fixed published score, *B* is known and the audit is particularly clean: there is no score-learning uncertainty. The adequacy question is target-population specific, however, because *α* and Σ can vary across cohorts even when the published weights do not. A score can therefore be adequate in one exposure–population combination and inadequate in another. This is not a defect of the score; it is a reminder that mediation adequacy is relational among the score, exposure mechanism, mediator covariance, and target population.

### 2.11 Interpretation of the diagnostics

The framework separates three questions that should not be collapsed. The exact/minimax geometry asks whether the retained subspace contains the whitened exposure direction and summarizes the worst-case loss by *ω*_*B*_. The benchmark geometry uses (*m*_*B*_, *γ*_*B*_) to describe relative distortion across an outcome-symmetric reference distribution. The observed-outcome geometry reduces to the scalar *D*_*B*_ once a particular outcome is introduced. A nonsignificant *D*_*B*_ therefore does not establish structural adequacy, and a large *ω*_*B*_ does not imply a particular realized error for one outcome.

The reallocation identity also gives the direct effect a simple interpretation. After residualizing on *W*, components of *M* omitted by *B*^*⊤*^*M* can remain associated with *A* and with *Y* ; their contribution is assigned to the coefficient on *A*. Consequently, direct effects from mediation models using different mediator summaries are different estimands, even when exposure, outcome, and covariates are unchanged. This is why total-effect stability cannot diagnose aggregation adequacy and why score selection should, when possible, be performed without using the downstream outcome.

### 2.12 Four analytic examples

Small examples make clear that the phenomenon is not a high-dimensional numerical artifact. Throughout, take standardized mediator residual variances for convenience.

#### Example 1

**when a simple mean really is exact**. Let *p* = 2,

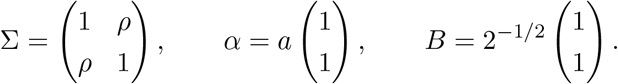

Then Σ^*−*1^*α* = *a*(1 + *ρ*)^*−*1^(1, 1)^*⊤*^ ∈ col(*B*), so the mean is exactly adequate for every *β* despite correlation. Symmetry, not independence, is what saves the mean. If the exposure effects become (*a*_1_, *a*_2_)^*⊤*^ with *a*_1_≠*a*_2_, the mean generally ceases to be exact because Σ^*−*1^*α* is no longer constant across components. This example illustrates why an annotation-defined mean can be defensible when both the exposure mechanism and covariance are approximately exchangeable within the group.

#### Example 2: an exposure-weighted score with zero systematic bias and nonzero fragility

Let *α* = (1, 0)^*⊤*^ and take the same Σ, but use *B* = *α* = *e*_1_. The raw exposure score retains the only mediator with a nonzero marginal exposure coefficient. Nevertheless

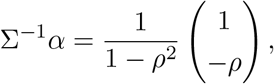

which is outside col(*B*) whenever *ρ*≠0. Direct calculation gives

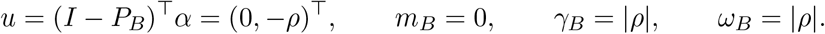

Thus the score has no systematic distortion under the spherical benchmark, but its unpredictable distortion grows one-for-one with residual correlation. At *ρ* = 0.8, half of benchmark outcome directions have relative gaps inside a Cauchy half-range of width 0.8 around zero, and the benchmark probability of a sign reversal is 1 − *F*_*B*_(1). The problem is not that the second mediator responds directly to the exposure; it does not. The problem is that conditional exposure information lives partly in that mediator because it is correlated with the first.

#### Example 3: the leading variance direction can retain none of the exposure direction

Let Σ = diag(4, 1), so the first mediator is the leading residual PC, and let *B* = *e*_1_. If *α* = (0, 1)^*⊤*^, all exposure signal lies in the second, lower-variance direction. Then *P*_*B*_ = *e*_1_*e*^*⊤*^_1_ and

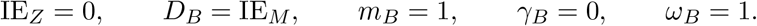

The leading PC explains 80% of residual mediator variance and exactly 0% of the exposure-relevant direction. Every mediated effect is assigned to the direct effect. This extreme case formalizes a familiar warning: variance explained is not the same criterion as causal information retained.

#### Example 4: exact compression does not improve overlap

For arbitrary *p*, take the one-dimensional exact score *B* = Σ^*−*1^*α*. Proposition 2 gives *D*_*B*_ = 0 for all *β*, so *m*_*B*_ = *γ*_*B*_ = *ω*_*B*_ = 0. Proposition 4, however, gives *r*_*Z*_ = *r*_*M*_ . The entire mediated-effect-relevant exposure direction can therefore be compressed to one scalar, but that compression does not restore positivity. Dimension and overlap are separate statistical notions: a one-dimensional covariate can predict exposure almost perfectly.

These examples are summarized in Table 2. They also suggest a useful diagnostic hierarchy. A mean can be exact under symmetry; an exposure score needs covariance alignment; a high-variance PC can be irrelevant to exposure; and the exact score can still be statistically difficult because it preserves all exposure predictability.

**Table 1.** Core notation used throughout the paper.

| Symbol | Meaning |
| --- | --- |
| $M \in \mathbb{R}^p, Z = B^\top M \in \mathbb{R}^k$ | full mediator vector and proposed aggregate(s) |
| $\alpha, \beta, \Sigma$ | exposure–mediator effects, mediator–outcome effects, residual mediator covariance |
| $P_B = B(B^\top \Sigma B)^{-1} B^\top \Sigma$ | oblique projection induced by the aggregation |
| $D_B = \text{IE}_M - \text{IE}_Z$ | realized aggregation gap |
| $\kappa = \alpha^\top \Sigma^{-1} \alpha$ | full exposure information in the mediators |
| $\kappa_B = \alpha^\top B(B^\top \Sigma B)^{-1} B^\top \alpha$ | exposure information retained by $B$ |
| $\omega_B^2 = 1 - \kappa_B / \kappa$ | distribution-free minimax adequacy loss |
| $(m_B, \gamma_B)$ | location and scale of the spherical Cauchy relative-gap profile |

**Table 2.**
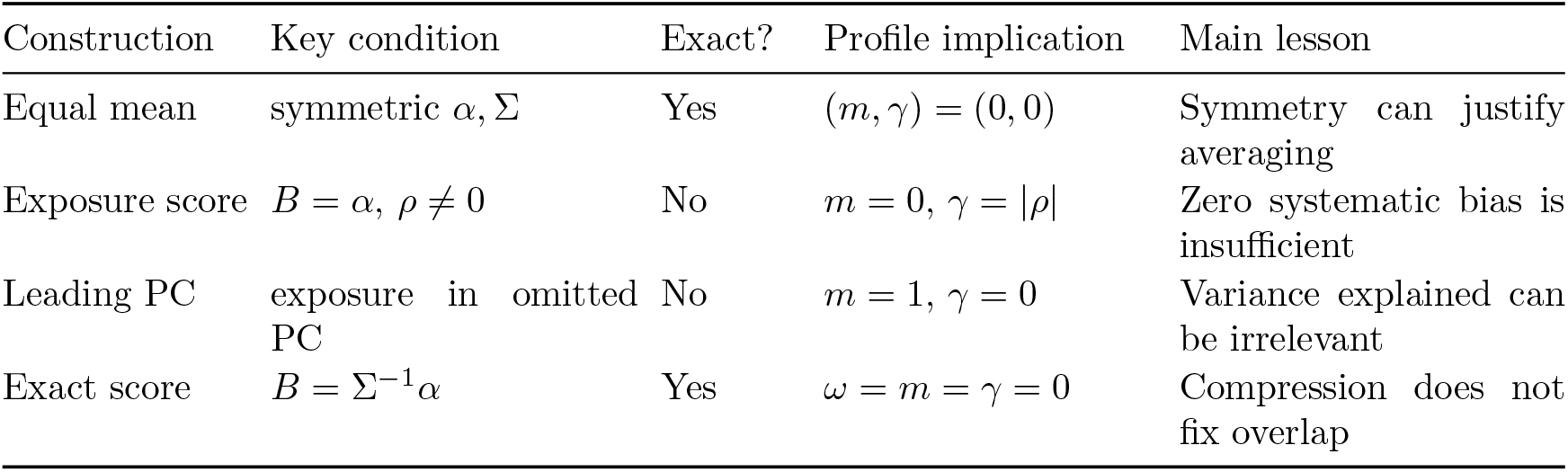
Analytic two-mediator archetypes. “Exact” means preservation for every outcome direction.

### 2.13 Aggregation adequacy is not causal identification

The results in this paper are deliberately conditional on the causal assumptions that justify interpreting the linear decomposition as mediation. Adequacy asks whether replacing *M* by *B*^*⊤*^*M* preserves that decomposition; it does not establish exchangeability, consistency, positivity, temporal ordering, or absence of exposure-induced mediator–outcome confounding. An exactly adequate score can preserve a biased mediation estimand if the underlying causal assumptions fail, and an inadequate score can be estimated with impeccable statistical precision.

This separation is useful because aggregation is often introduced late in an analysis as a computational convenience. Suppose the full mediator model would identify IE_*M*_ under a stated causal graph, but the analyst substitutes a pathway score to reduce dimension. The causal graph alone does not imply that the score-level effect equals the component-level effect. Proposition 2 adds the missing statistical condition. Conversely, if the scientific estimand is explicitly defined as mediation through a published score—for example an epigenetic clock itself—then no component-level preservation claim is needed. The score is the mediator of substantive interest, and the audit answers a different question: whether statements about the score can also be extended to the CpGs from which it was constructed.

Positivity deserves special attention because it can be obscured by dimension reduction. Standard diagnostics performed only after replacing *M* by *Z* may show reasonable residual exposure variation and suggest that the aggregate model is well behaved. Equation (5) shows exactly what was purchased by that improvement: exposure-predictive information was discarded. If the scientific claim concerns the full mediator set, better overlap in the aggregate model is not free evidence of better identification; it can reflect movement to a different estimand.

Temporal ordering is equally separate. The TCGA gene-expression illustration has methylation and expression measured from the same tumour specimen. The aggregation audit is still alge-braically informative because it compares two decompositions under the same working temporal assumption, but it cannot validate that assumption. A prospective cohort with exposure measured first, methylation second, and expression or clinical outcome later would strengthen causal interpretation without changing the adequacy mathematics. For this reason we treat the application as an estimand illustration rather than a biological claim.

Finally, the framework does not imply that component-level mediation is always the preferable target. Biological systems are often intentionally represented by latent or composite constructs. The methodological requirement is narrower: name the target correctly. If the target is the composite, analyze the composite. If the target is mediation through all measured components, demonstrate or quantify how the composite relates to that target rather than assuming equivalence from dimensionality reduction alone.

## 3 An outcome-free adequacy profile

### 3.1 The Cauchy profile

Proposition 2 is a knife-edge characterization; most practical aggregates fail it. To quantify *how badly* an aggregate fails without committing to an outcome model, we average over outcome effects.

#### Theorem 1

(Spherical Cauchy adequacy profile). *Fix α*≠0, Σ, *and B, and let u* = (*I* − *P*_*B*_)^*⊤*^*α. Suppose the outcome-effect vector β has a non-degenerate spherically symmetric distribution about zero, so β* = *RU with R >* 0 *and U uniform on the unit sphere. Then, whenever γ*_*B*_ *>* 0,

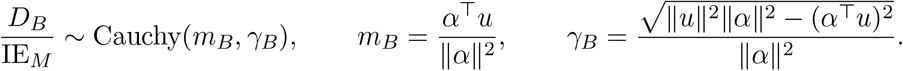

*If γ*_*B*_ = 0 *the ratio equals m*_*B*_ *almost surely. The result therefore includes Gaussian outcome effects, multivariate t effects, and arbitrary radial scale mixtures as special cases*.

The theorem follows because the radial component cancels in the ratio and a uniform direction can be represented as *G/*∥*G*∥ with *G* ∼ *N* (0, *I*_*p*_). Component ratios of spherical vectors inherit the Cauchy law (Arnold and Brockett, 1992). We call (*m*_*B*_, *γ*_*B*_) the *adequacy profile* of *B*. The location *m*_*B*_ = 1 − *α*^*⊤*^*P*_*B*_*α/*∥*α*∥_2_ is the regression of the gap on the true indirect effect under the spherical benchmark. The scale *γ*_*B*_ measures outcome-to-outcome unpredictability; for half of benchmark outcome directions the relative gap lies in *m*_*B*_ ± *γ*_*B*_. Both quantities involve only (*A, W, M*) and the proposed *B*.

#### Corollary 2

*(i) m*_*B*_ = 0 *whenever α* ∈ col(*B*). *(ii) m*_*B*_ = *γ*_*B*_ = 0 *if and only if* Σ^*−*1^*α* ∈ col(*B*). *(iii) If the columns of B are orthonormal eigenvectors of* Σ, *then P*_*B*_ = *BB*^*⊤*^, *m*_*B*_ = ∥(*I* − *BB*^*⊤*^)*α*∥^2^*/*∥*α*∥^2^ ∈ [0, 1], *and* 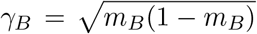 *the profile depends on* Σ *only through the retained eigenspace*.

Part (i) explains why an exposure-signature score (*B* ∝ *α*) can have zero systematic distortion while remaining unreliable when *γ*_*B*_ is large. Part (iii) shows that principal-component subspaces can only understate mediation systematically under this benchmark. The identity 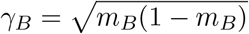 is exact only when the retained directions are eigenvectors of the residual covariance Σ; empirical PCs computed from a different covariance matrix need not satisfy it.

### 3.2 From a profile to error probabilities

A location–scale pair is useful diagnostically, but investigators often want a probability on a scientifically meaningful scale. Let *R*_*B*_ = *D*_*B*_*/*IE_*M*_ and, for *γ*_*B*_ *>* 0, write

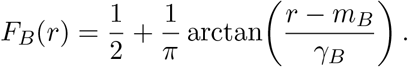

For any tolerance *c >* 0, the benchmark probability that aggregation changes a nonzero mediated effect by more than a fraction *c* is

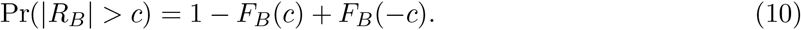

Because IE_*Z*_ = IE_*M*_ (1 − *R*_*B*_), the benchmark sign-reversal probability is

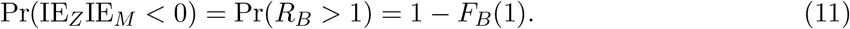

The regions *R*_*B*_ *<* 0, 0 *< R*_*B*_ *<* 1, and *R*_*B*_ *>* 1 correspond respectively to same-sign overstatement, same-sign attenuation, and sign reversal. These probabilities are not posterior probabilities about any particular biological outcome; they summarize the geometry of a proposed aggregation under an outcome-symmetric reference distribution. They are therefore best interpreted as pre-outcome stress tests.

#### Remark 2

(Choice of benchmark). Spherical symmetry expresses ignorance about which standardized mediator components matter for the outcome. It is not a biological prior. If investigators have a scientifically defensible anisotropic benchmark, the same projection identity can be integrated under that distribution numerically. Placing spherical symmetry on Σ^1*/*2^*β* instead of *β* yields a whitened benchmark aligned with the minimax geometry in Section 2.6. The sparse simulations in Section 5 examine robustness to a strongly non-spherical departure.

### 3.3 Estimation

The profile depends on *α*, on Σ only through the *k* × *k* matrix *B*^*⊤*^Σ*B* and the products Σ*B*, and on inner products of *α* with itself and with *B*. It can therefore be estimated when *p* ≫ *n* without inverting Σ. Plug-in estimation is biased because 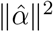 and the quadratic forms in 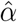 absorb sampling noise. We use sample splitting.

Partition the sample into four folds, stratified by exposure. In fold *j*, regress *M* on (1, *A, W*) to obtain 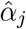 and the residual covariance *S*_*j*_; let 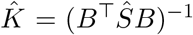 be computed from the full sample. Under Gaussian errors, 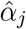 sis independent of *S*_*j*_ and of all quantities from other folds, so cross-products formed from distinct folds are unbiased for their population counterparts. Writing 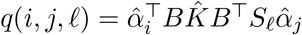, we average over the 24 ordered quadruples (*i, j, k, ℓ*) of distinct folds:

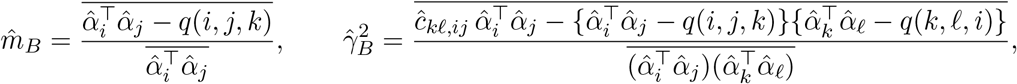

where 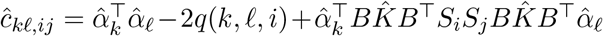 estimates ∥*u*∥^2^, and bars denote averages over quadruples. Using four folds ensures that each factor of the squared term in 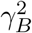is built from independent data; a two-fold version, which reuses the same pair of coefficient vectors, is biased downward (Section 5.5). We report 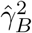 without truncation, which is appropriate for testing *γ*_*B*_ = 0.

The shared full-sample 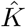 is a remaining source of finite-sample bias that grows with *k*; Section 5.5 maps the range in which the estimator is reliable.

Sampling uncertainty in the profile should be reported rather than treating 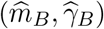 as fixed diagnostics. We use an exposure-stratified nonparametric bootstrap that repeats the fold assignment and, for learned scores, refits *B* inside each resample. Percentile intervals can be reported for *m*_*B*_ and for 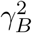 ; because 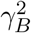 is on the boundary under exact adequacy, we retain its signed cross-fitted estimate for inference and use 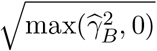 only for descriptive plots. This bootstrap also propagates the uncertainty in 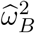 from Section 2.7 when the full information estimator is available.

### 3.4 Fixed and data-learned aggregations

The algebra above treats *B* as fixed. This is literal for published clock weights, externally supplied pathway weights, and prespecified gene averages, but not for principal components or sign scores learned in the analysis sample. Treating a learned 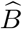 as fixed can understate uncertainty and can entangle score construction with outcome analysis.

A simple honest protocol is to separate construction from evaluation. Split the sample into training and evaluation observations; learn 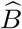 using only (*A, W, M*) in the training sample; freeze those weights; and compute the profile, *ω*_*B*_, and any outcome analysis in the evaluation sample. Conditional on the training sample, all fixed-*B* results apply exactly to 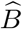. Repeating this operation across folds can recover efficiency, but the estimand must be stated carefully: a cross-fitted analysis averages the effects of fold-specific learned scores, whereas a final score refitted on the entire sample is a different random object. Nested bootstrap or repeated sample splitting is preferable when inference is desired for the full learning algorithm.

This distinction matters empirically. In Section 7, the sign and PC weights are re-estimated in each bootstrap resample; their intervals are therefore wider than intervals that would condition on the observed weights. For scientific scores intended for transport to other cohorts, the cleanest design is to estimate and freeze the score in an external training sample and audit it in an independent target sample.

## 4 Inference for the realized aggregation gap

The outcome-free diagnostics answer whether a score can fail and how severely it can fail across outcome directions. Once a particular *Y* is observed, a different question becomes available: is the realized gap *D*_*B*_ detectably different from zero? Proposition 1 gives an unusually simple route because

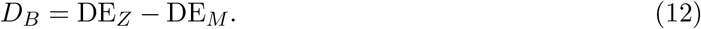

Thus inference can be based on the difference between two direct-effect regressions, provided their strong sampling dependence is retained.

### 4.1 Paired influence-function inference

For *s* ∈ {*M, Z*}, let *X*_*M*_ = (*W*^*⊤*^, *M*^*⊤*^)^*⊤*^ and *X*_*Z*_ = (*W*^*⊤*^, *Z*^*⊤*^)^*⊤*^, let *θ*_*s*_ denote the coefficient of *A* in the population linear projection of *Y* on (1, *A, X*_*s*_), and define

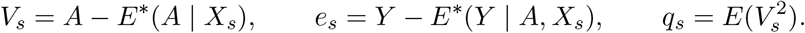

Then *θ*_*M*_ = DE_*M*_, *θ*_*Z*_ = DE_*Z*_, and *D*_*B*_ = *θ*_*Z*_ − *θ*_*M*_ .

#### Theorem 2

(Paired inference for *D*_*B*_). *Assume iid observations, fixed dimensions, finite fourth moments, positive q*_*M*_, *q*_*Z*_, *and nonsingular population Gram matrices for the two regressions. Let* 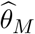*and* 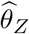 *be ordinary least-squares coefficients of A and* 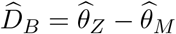. *Then*

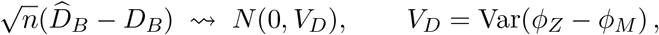

*where*

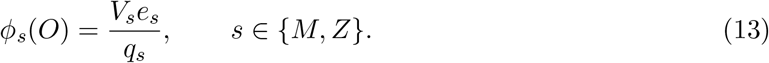

*A consistent heteroskedasticity-robust variance estimator is the sample variance of ϕ*_*Zi*_ − *ϕ*_*Mi*_, *divided by n*.

The theorem yields the Wald interval

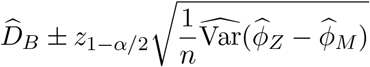

and a direct test of *H*_0_ : *D*_*B*_ = 0. The covariance term is essential. Computing 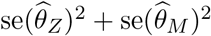] as if the two fits were independent discards the fact that they use the same observations, exposure, outcome, and much of the same adjustment information. In the simulation of Section 6, that independence calculation is roughly 2.2 to 3.6 times wider under the null.

The test is outcome-specific. Failure to reject *D*_*B*_ = 0 does not establish that the aggregation is universally adequate: cancellation between the omitted exposure and outcome directions can make the realized gap zero even when *ω*_*B*_, *m*_*B*_, or *γ*_*B*_ is large. Conversely, a significant *D*_*B*_ is direct evidence that the aggregate and the full mediator vector answer different mediation questions for the observed outcome.

### 4.2 High-dimensional full mediator models: scope of the present theory

Theorem 2 is a fixed-dimensional result. When *p* is large enough that the full regression cannot be fit by ordinary least squares, Equation (12) still suggests a useful route, but this paper does not claim a high-dimensional gap theorem. One possible extension starts from partially linear representations

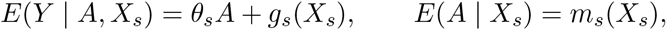

for *s* ∈ {*M, Z*} and uses cross-fitted orthogonal scores of the double/debiased-machine-learning type (Chernozhukov et al., 2018). Establishing root-*n* inference for the paired difference requires explicit product-rate, overlap, and conditional-mean conditions for both adjustment sets and care when the aggregate is itself learned. Those conditions are stronger than the projection assumptions used for Proposition 1; we therefore leave a formal high-dimensional theorem to future work.

This limitation does not affect the moderate-dimensional theorem or the outcome-free audit. In particular, gene-level *ω*_*B*_ estimation can remain feasible when the genome-wide mediator vector has *p* ≫ *n*, because each gene contains only a modest number of CpGs. Cross-fitting can protect high-dimensional nuisance estimation from overfitting, but it cannot repair the positivity problem identified by Proposition 4.

### 4.3 Recommended inferential workflow

The preceding results suggest a staged analysis. Before examining *Y*, report the dimension *k*, the retained exposure-information fraction 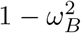 using Section 2.7 when estimable, and the Cauchy profile (*m*_*B*_, *γ*_*B*_) with bootstrap uncertainty and threshold probabilities such as (10) and (11). After introducing *Y*, estimate both IE_*M*_ and IE_*Z*_ when feasible and report 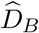 with a paired interval. For learned scores, construct weights out of fold and either condition on the training split or bootstrap the complete learning procedure. Finally, interpret a score-level indirect effect as mediation through that score, not automatically as mediation through all components that were used to construct it. This ordering also reduces researcher degrees of freedom. An aggregation can be rejected as structurally inadequate using only (*A, W, M*), before the outcome effect is known. Candidate scores can therefore be compared without selecting the one that happens to produce the most favorable mediation result.

## 5 Semi-synthetic evaluation

### 5.1 Data and population

We used Illumina 450K methylation for primary lung adenocarcinomas from The Cancer Genome Atlas, obtained from UCSC Xena (Goldman et al., 2020; Wang and Liu, 2019). After restricting to tumours with smoking history, age, sex, pathologic stage, and consensus purity (Aran et al., 2015), 421 tumours remained (62 lifelong never-smokers, 359 ever-smokers). Probe filtering removed probes with more than 10% missingness, near-zero variance, sex-chromosome location, or known SNPs at the probe, CpG, or extension base, leaving 285,755 CpGs, which were converted to M-values. Regressing each CpG on smoking, age, sex, stage, and purity with limma (Ritchie et al., 2015) identified 4,625 CpGs at false discovery rate 0.05; these standardized CpGs define the mediator vector (*p* = 4,625).

The population used in all simulations takes *α* to be the estimated smoking coefficients and 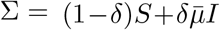, where *S* is the residual covariance of the standardized CpGs, 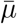 its average eigenvalue, and *δ* = 0.2 unless stated otherwise (Ledoit and Wolf, 2004). Shrinkage is necessary: with *p > n* the unshrunk residual covariance is singular, the full-mediator indirect effect is not identified in the empirical population, and exposure-discriminant scores computed from it degenerate. All population quantities below are exact closed-form evaluations of Proposition 1 and Theorem 1 computed through the singular value decomposition of *S*, so no *p* × *p* matrix is formed.

We compared the following aggregations: a sign score (mean of hypermethylated minus mean of hypomethylated CpGs, mimicking direction-weighted pathway activation); separate hyper- and hypomethylation means (*k* = 2); an exposure-weighted score (*B* = *α*); leading principal components of the CpG matrix (*k* ∈ {1, 2, 5, 10, 20, 50, 100}); and the exact score Σ^*−*1^*α* as a numerical check.

### 5.2 Adequacy of common aggregations

Table 3 reports the adequacy profile of each aggregation and summaries over 1,000 random outcome models in which 50 CpGs, drawn at random, had Gaussian effects. Among draws with an above-median true indirect effect, the sign score misstated the effect by more than 50% in 19% of draws and PC1 did so in 34%. Among the 20% of draws with the smallest true indirect effect, the sign score reported an aggregate effect larger than the median true effect in 12% of draws (*false mediation*); the exposure-weighted score did so in 18.5%. Sign reversal among non-trivial effects was rare (0–7%): aggregation distorts magnitude far more often than direction.

**Table 3.** Adequacy of common aggregations in the TCGA semi-synthetic population (*δ* = 0.2). Profile values are exact. Random-draw summaries use 1,000 outcome models with 50 randomly placed Gaussian effects; “non-trivial” denotes draws with |IE_*M*_ | above the median, “weak” the lowest quintile.

| Aggregation | Profile |  | Non-trivial effects |  |  | Weak effects |
| --- | --- | --- | --- | --- | --- | --- |
| | $m_B$ | $\gamma_B$ | median<br>$\vert D_B/\text{IE}_M\vert $ | % error<br>> 50% | % sign<br>flip | % false<br>mediation |
| Sign score | −0.068 | 0.412 | 0.249 | 19.0 | 1.0 | 12.0 |
| Hyper/hypo means | 0.006 | 0.327 | 0.183 | 9.4 | 0.2 | 5.5 |
| Exposure-weighted | 0.000 | 0.503 | 0.289 | 28.0 | 2.8 | 18.5 |
| PC1 | 0.283 | 0.481 | 0.353 | 34.4 | 7.4 | 15.5 |
| PC2 | 0.119 | 0.327 | 0.206 | 10.8 | 0.2 | 5.5 |
| PC5 | 0.062 | 0.271 | 0.166 | 7.4 | 0.0 | 3.5 |
| PC10 | 0.049 | 0.247 | 0.142 | 5.4 | 0.2 | 2.0 |
| PC20 | 0.043 | 0.236 | 0.142 | 4.2 | 0.2 | 2.0 |
| PC50 | 0.031 | 0.218 | 0.130 | 2.8 | 0.2 | 2.5 |
| PC100 | 0.022 | 0.208 | 0.118 | 2.6 | 0.0 | 0.5 |
| $\Sigma^{-1}\alpha$ (exact) | 0.000 | 0.000 | 0.000 | 0.0 | 0.0 | 0.0 |

Weighting CpGs by their exposure effects was worse than using signs alone (Table 3; Figure 1a). This is predicted by Corollary 2(i): *B* ∝ *α* eliminates *m*_*B*_ but leaves *γ*_*B*_ unconstrained, and weighting by *α* concentrates the score on the most strongly correlated smoking-responsive CpGs. The correct weights are Σ^*−*1^*α*, not *α*. Principal-component scores improved quickly up to about ten components and then plateaued (*γ*_*B*_*≈*0.21 at *k* = 100), because variance-ordered directions do not target the direction that matters.

**Figure 1.**
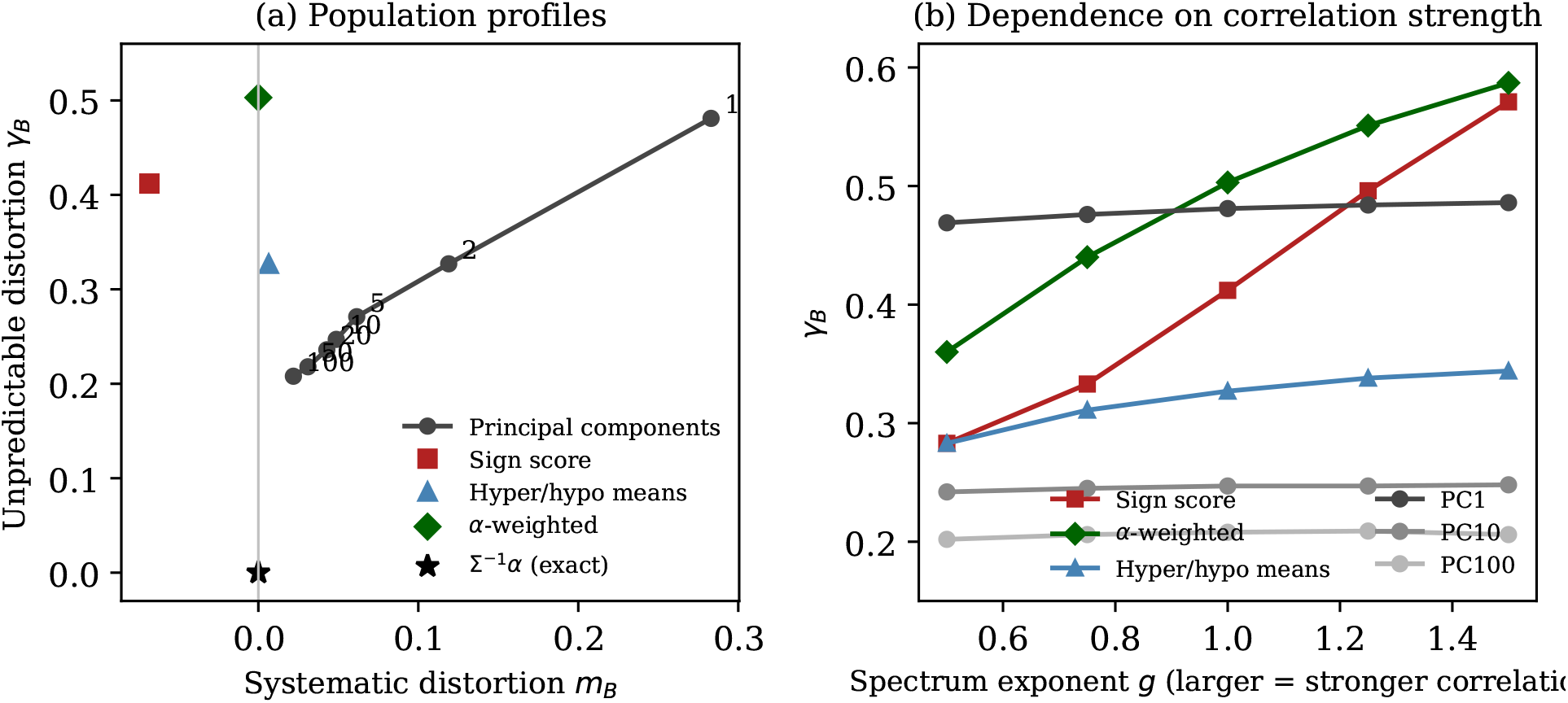
(a) Adequacy profiles (*m*_*B*_, *γ*_*B*_) in the TCGA semi-synthetic population; numbers label the count of principal components. (b) Unpredictable distortion *γ*_*B*_ as the eigenvalue spectrum of the residual covariance is raised to the power *g* and rescaled to constant total variance. *Alt text:* Two-panel diagnostic plot. Panel (a) places common aggregation methods in the plane of systematic distortion *m*_*B*_ and unpredictable distortion *γ*_*B*_; exact aggregation is at the origin and principal-component labels show the number retained. Panel (b) plots *γ*_*B*_ against increasing residual-correlation concentration, with sign and exposure-weighted scores becoming more fragile while principal-component profiles change little.

### 5.3 Accuracy of the Cauchy approximation

For each aggregation, the median and half-interquartile range of *D*_*B*_*/*IE_*M*_ over 2,000 dense Gaussian outcome models reproduced *m*_*B*_ and *γ*_*B*_ to about two decimals (for example PC1: 0.284 and 0.484 against 0.283 and 0.481; sign score: −0.056 and 0.413 against −0.068 and 0.412). The same summaries from the sparse draws of Table 3 were similarly close (PC1: 0.274 and 0.481; sign score: −0.078 and 0.439), confirming that the profile is informative when only a few mediators affect the outcome. Consistent with the theorem, the summaries were insensitive to sparsity (*s* ∈ {10, 50, 200}) and to shrinkage (*δ* ∈ {0.1, 0.2, 0.5}): across this grid the percentage of non-trivial effects misstated by more than half ranged from 18.2 to 20.8 for the sign score and from 33.2 to 35.4 for PC1.

### 5.4 Dependence on correlation strength

To vary the strength of mediator correlation, we replaced the eigenvalues *d*_*j*_ of *S* by 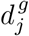, rescaled to preserve total variance. Figure 1b shows that *γ*_*B*_ for the sign score rose from 0.28 to 0.57 and for the exposure-weighted score from 0.36 to 0.59 as *g* increased from 0.5 to 1.5, while principal-component profiles were essentially unchanged, as Corollary 2(iii) predicts. The systematic distortion of the sign score changed sign across this range (*m*_*B*_ from 0.02 to −0.14): whether a direction-weighted pathway score overstates or understates mediation depends on a correlation structure that the analyst does not observe.

### 5.5 Estimating the profile and building scores from data

We drew samples of *n* = 421 from the population, with exposure and covariates fixed at their observed values, and applied the four-fold estimator (100 replicates). Estimates of *m*_*B*_ were close to the truth for every aggregation (Table 4). Estimates of *γ*_*B*_ were within about 10% of the truth for *k* ≤ 20 (0.220 against 0.236 at *k* = 20); at *k* = 50 they were biased downward (0.156 against 0.218) and at *k* = 100 the estimator failed, with 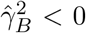 in 65% of replicates. We therefore recommend the *γ*_*B*_ estimator for *k* ≲ *n/*20. A two-fold version was biased downward by 3–13% even for *k* = 1, and a naive inverse-Wishart correction to 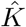 destroyed the accuracy of 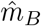. For the exact score,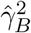 was centred near zero but highly variable (standard deviation 0.60), reflecting the difficulty of estimating low-variance directions.

**Table 4.** Four-fold estimation of the adequacy profile (*n* = 421, 100 replicates) and true profiles of scores rebuilt in each sample (30 replicates). Half-range: interval containing the relative gap for half of outcome models, *m*_*B*_ ± *γ*_*B*_.

**Panel A. Estimating a fixed aggregation’s profile**
| Aggregation | $\hat{m}_B$ | $m_B$ | $\hat{\gamma}_B$ | $\gamma_B$ |
| --- | --- | --- | --- | --- |
| Sign score | −0.061 | −0.068 | 0.423 | 0.412 |
| Hyper/hypo means | 0.009 | 0.006 | 0.334 | 0.327 |
| Exposure-weighted | 0.002 | 0.000 | 0.510 | 0.503 |
| PC1 | 0.280 | 0.283 | 0.482 | 0.481 |
| PC10 | 0.051 | 0.049 | 0.240 | 0.247 |
| PC20 | 0.045 | 0.043 | 0.220 | 0.236 |
| PC50 | 0.031 | 0.031 | 0.156 | 0.218 |
| PC100 | 0.018 | 0.022 | fails | 0.208 |

**Panel B. True profiles of scores rebuilt from each sample**
| Score | $m_B$ | $\gamma_B$ | Half-range |
| --- | --- | --- | --- |
| PC50 | 0.031 | 0.226 | −0.20 to 0.26 |
| PC20 | 0.041 | 0.240 | −0.20 to 0.28 |
| PC10 | 0.047 | 0.248 | −0.20 to 0.30 |
| Hyper/hypo means | 0.006 | 0.327 | −0.32 to 0.33 |
| Lasso, $A$ on $M$ | 0.092 | 0.341 | −0.25 to 0.43 |
| Elastic net, $A$ on $M$ | 0.067 | 0.350 | −0.28 to 0.42 |
| Sign score | −0.068 | 0.412 | −0.48 to 0.34 |
| PC1 | 0.277 | 0.482 | −0.21 to 0.76 |
| Exposure-weighted | 0.008 | 0.502 | −0.49 to 0.51 |

We also rebuilt each practitioner score from each sample and evaluated its true population profile (30 replicates). Sample-based principal components performed essentially as in the population (*γ*_*B*_ between 0.23 and 0.25 for *k* from 10 to 50). Lasso and elastic-net regressions of exposure on the mediators, the sample analogues of the exact score (Proposition 4), achieved *γ*_*B*_*≈*0.34–0.35 with small average *m*_*B*_ but large replicate-to-replicate variability (standard deviation of *m*_*B*_ 0.17– 0.19). In a smaller exploratory run (three replicates), ridge-regularized discriminant scores were systematically biased over most of the penalty path (*m*_*B*_ between −0.25 and −0.40), approaching the exposure-weighted score only under very heavy penalization. At *p/n≈*11, the exactly adequate score exists in theory but cannot be recovered from data; the practical value of the theory lies in auditing proposed scores.

## 6 Finite-sample study of gap inference

We evaluated Theorem 2 in a separate simulation whose sole purpose is inference for *D*_*B*_. This study does not reuse the TCGA empirical population and therefore provides a clean check of the asymptotic argument. We generated two Gaussian covariates, a continuous exposure with residual variance one, and *p* ∈ {12, 30} mediators from model (1) with an AR(1) residual covariance (*ρ* = 0.35). The candidate aggregation was the standardized equal-weight mean. Under the null, *β* was chosen in col(*B*) so that *D*_*B*_ = 0 exactly and IE_*M*_ = 0.35. Under the alternative, a component in the omitted gap direction was added so that *D*_*B*_ = 0.20 exactly. The outcome error variance was one. We used 800 Monte Carlo replicates for each setting.

Table 5 compares the paired influence-function interval with a deliberately naive interval that treats 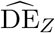 and 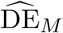 as independent. In the native-R simulation, paired coverage under the null ranged from 95.8% to 97.5%, and across all eight null/alternative cells it ranged from 93.3% to 97.5%. The empirical standard deviation closely tracked the mean estimated standard error. Under *D*_*B*_ = 0.20, rejection was at least 99.9% in every setting shown. Under the null, the independence interval was about 2.2 to 3.6 times wider and covered 100% of the time; at *n* = 300, *p* = 12 its rejection rate under the alternative fell to 68.3%. Thus the paired covariance term materially improves both interval precision and power.

**Table 5.** Finite-sample inference for the aggregation gap. Paired intervals use Theorem 2; naive intervals ignore covariance between the two direct-effect estimates. Each row uses 800 replicates.

| $n$ | $p$ | Scenario | Bias | Emp. SD | Mean SE | Coverage | Reject $H_0$ | Width (naive) |
| --- | --- | --- | --- | --- | --- | --- | --- | --- |
| 300 | 12 | $D_B = 0$ | 0.001 | 0.025 | 0.026 | 0.975 | 0.025 | 0.103 (0.332) |
| 300 | 12 | $D_B = 0.20$ | 0.001 | 0.044 | 0.043 | 0.939 | 0.999 | 0.169 (0.357) |
| 600 | 12 | $D_B = 0$ | -0.000 | 0.016 | 0.017 | 0.958 | 0.043 | 0.065 (0.235) |
| 600 | 12 | $D_B = 0.20$ | 0.000 | 0.028 | 0.029 | 0.956 | 1.000 | 0.115 (0.253) |
| 600 | 30 | $D_B = 0$ | -0.001 | 0.028 | 0.029 | 0.960 | 0.040 | 0.113 (0.250) |
| 600 | 30 | $D_B = 0.20$ | 0.000 | 0.030 | 0.032 | 0.968 | 1.000 | 0.125 (0.257) |
| 1000 | 30 | $D_B = 0$ | -0.000 | 0.020 | 0.021 | 0.965 | 0.035 | 0.084 (0.194) |
| 1000 | 30 | $D_B = 0.20$ | 0.000 | 0.025 | 0.024 | 0.933 | 1.000 | 0.094 (0.199) |

The last alternative cell shows mild undercoverage (93.3%), which is useful rather than embarrassing: the first-order sandwich approximation is not exact even at *n* = 1000. The *n* = 300, *p* = 12 alternative cell also shows modest undercoverage (93.9%). A bootstrap of the paired estimat-ing equations or a small-sample robust correction may improve this behavior, and we leave that comparison for future work. The purpose of the study is narrower: the influence-function variance tracks Monte Carlo variability well and materially improves on an independence calculation. The complete simulation, including the exact population construction, is reproduced in base R by `03_gap_inference_simulation.R`; its exact table source is `gap_inference_results_R.csv`.

## 7 Application: an outcome-free genomic audit and an expression illustration

### 7.1 A cautionary first attempt

High-dimensional mediation methods have frequently been illustrated with smoking, tumour methylation, and survival in TCGA lung cancer (Luo et al., 2020). We first attempted this design, using restricted mean survival time to three years via pseudo-observations (Andersen et al., 2004) as the outcome. After adjustment for age, sex, stage, and purity the total effect of smoking was −0.055 years (standard error 0.13; *p* = 0.68), so the survival analysis provided little information about a smoking effect. With the 4,625 screened CpGs as mediators, a cross-fitted partially linear estimate of DE_*M*_ (Chernozhukov et al., 2018) had standard error 0.17. The exposure nuisance model in this exploratory calculation was a Gaussian penalized regression of the binary smoking indicator, used only for partialling-out rather than as a propensity model; its fitted values therefore should not be interpreted as probabilities. Given the weak total-effect and direct-effect information, we did not pursue a survival mediation decomposition.

### 7.2 Genome-wide outcome-free audit

We next audited aggregation without using survival, gene expression, or any other downstream outcome. CpGs were mapped to genes using the UCSC gene annotation carried by the 450K array. Within each gene, CpG-level smoking *p*-values from the stage-1 exposure model were combined by the Simes procedure, and the resulting gene-level values were adjusted by Benjamini–Hochberg. We restricted attention to genes with 5–150 annotated CpGs and retained 1,314 genes at false discovery rate 0.05. For each gene, we estimated *κ* and *κ*_*B*_ using Proposition 6 with 25 exposure-stratified two-fold splits, adjusting for age, sex, stage, and purity; split-specific information estimates were averaged before forming 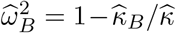 . No outcome variable entered screening, score construction,or evaluation.

The primary scores matched the scale of the detailed application: the mean of the original methylation M-values, the promoter mean when promoter probes were present, and a two-dimensional promoter-plus-non-promoter summary when both annotation classes were present. Estimates were deliberately left untruncated. This matters because the cross-fitted unbiased estimator of retained exposure information can be slightly negative in finite samples even though the population quantity is non-negative.

Because gene-level ratios become unstable when 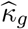 is close to zero, our primary collection-level summary is the *pooled ratio-of-sums loss*

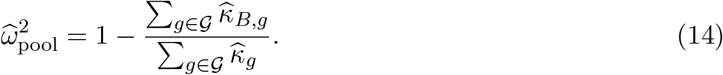

The numerator and denominator are sums of unbiased split-sample information estimators under Proposition 6; their ratio is not itself exactly unbiased, but it is substantially more stable than taking a median of ratios whose denominators can be near zero. We compare it with the *κ*-weighted random-subspace benchmark

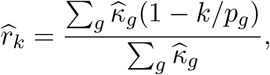

where *k* = 1 for a one-dimensional mean and *k* = 2 for the promoter-plus-non-promoter summary. Descriptive 95% intervals resample genes and recompute the ratio of sums; because genes share participants, these intervals describe stability of the empirical gene collection rather than participant-level sampling uncertainty.

In the full-data descriptive audit, the pooled gene-mean loss was 0.811 (95% gene-bootstrap interval 0.797–0.823), compared with a 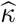-weighted one-dimensional random benchmark of 0.913 (Table 6). Thus the gene mean retained about 18.9% of the estimated exposure information versus 8.7% expected for the dimension-only benchmark, approximately 2.17 times as much. The promoter mean gave a similar pooled loss of 0.800. Retaining promoter and non-promoter means as two components improved the pooled loss to 0.625 (0.606–0.642), compared with a two-dimensional weighted random benchmark of 0.832. The improvement is therefore substantive but incomplete.

**Table 6.** Pooled outcome-free audit. The pooled loss is 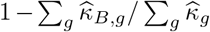. Intervals are descriptive 2,000-resample gene-level bootstrap intervals. The random benchmark weights 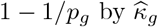. Cross-screen rows rank genes in one half and estimate adequacy only in the other half. The two-component column retains promoter and non-promoter means separately.

| Analysis | Gene set | median $\hat{\kappa}$ | % $\hat{\kappa} < 0.1$ | pooled gene-mean loss (95% interval) | weighted random loss | pooled two-component loss |
| --- | --- | --- | --- | --- | --- | --- |
| Full data | 1,314 FDR-selected genes | 0.371 | 0.4 | 0.811 (0.797, 0.823) | 0.913 | 0.625 |
| Screen 2 → estimate 1 | top 100 | 0.311 | 15.0 | 0.827 (0.762, 0.886) | 0.913 | 0.634 |
| Screen 2 → estimate 1 | top 250 | 0.256 | 24.8 | 0.836 (0.795, 0.875) | 0.914 | 0.634 |
| Screen 1 → estimate 2 | top 100 | 0.147 | 48.0 | 0.907 (0.842, 0.969) | 0.932 | 0.689 |
| Screen 1 → estimate 2 | top 250 | 0.155 | 42.8 | 0.932 (0.894, 0.969) | 0.927 | 0.714 |

The dimension comparison is clearest after stratifying by the number of CpGs (Figure 2). Gene-mean pooled loss rose from 0.702 for genes with 5–10 CpGs to 0.949 for genes with 61–150 CpGs, while the corresponding one-dimensional random benchmark rose from 0.858 to 0.987. The two-component summary was better in every bin, with pooled loss from 0.487 to 0.845. These results sharpen the interpretation of the earlier per-gene medians: a large absolute loss is partly inevitable when many mediator dimensions are compressed to one number, but annotation-based means recover only part of what is available beyond that dimension-only baseline.

**Figure 2.**
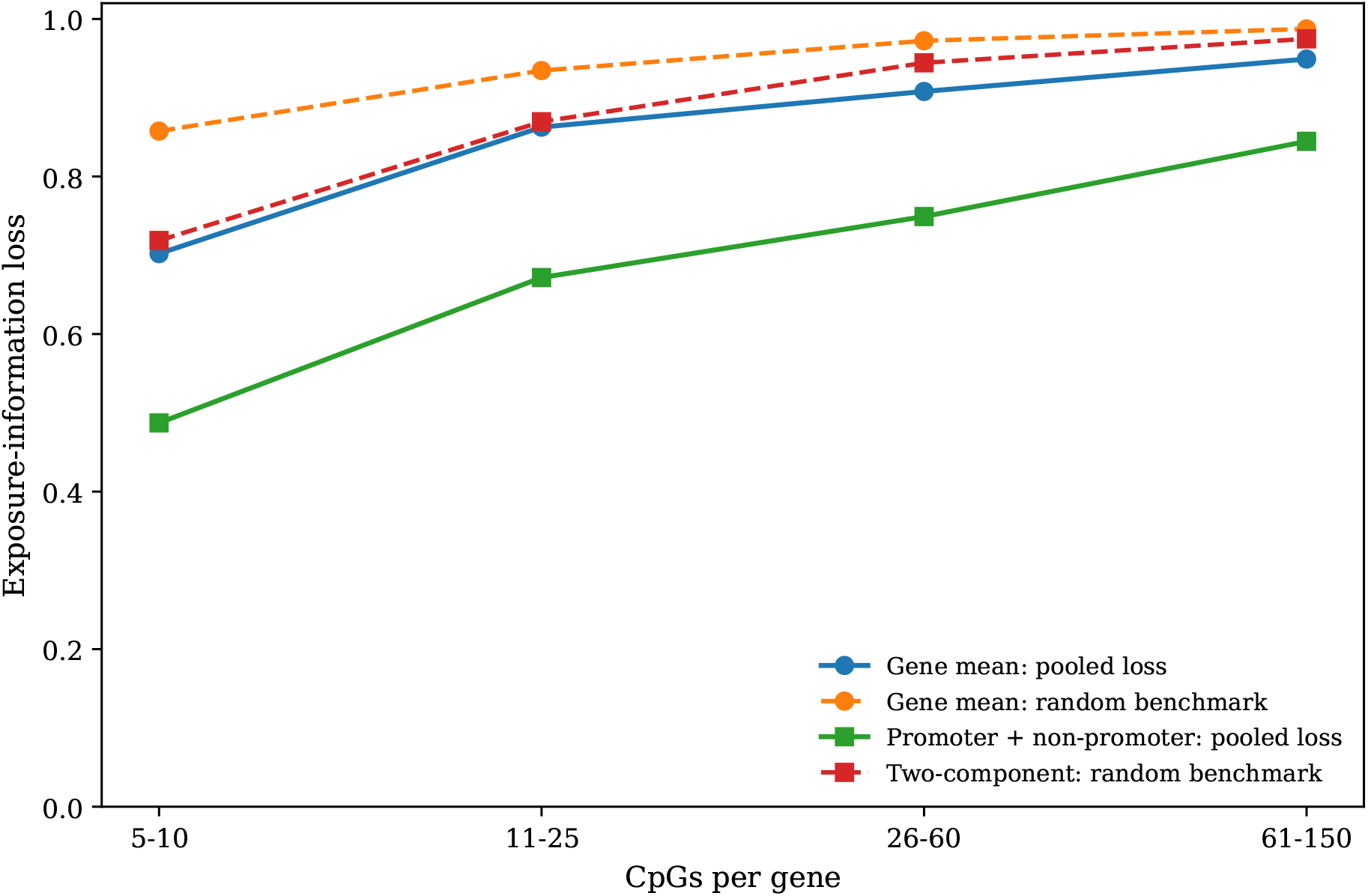
Full-data pooled exposure-information loss by gene size. Solid curves show the pooled loss of the annotation-based summaries; dashed curves show the corresponding *κ*-weighted random-subspace benchmarks 1 − *k/p*. The figure is descriptive because screening and estimation use the same participants.

As secondary descriptive diagnostics, the full-data per-gene gene-mean loss had median 0.879 (interquartile range 0.591–0.996), and the promoter-plus-non-promoter summary had median 0.601 (0.312–0.863). Gene-mean loss increased with the number of CpGs (Spearman *ρ* = 0.31) and decreased with mean absolute within-gene CpG correlation (*ρ* = −0.44). These individual ratios are not used as the primary evidence because 358 of 1,314 full-data gene-mean estimates (27.2%) fell outside [0, 1], a finite-sample consequence of unconstrained unbiased quadratic-form estimation. The full-data screen and the adequacy estimator use the same participants, so selection on smoking association can inflate estimated exposure information. The full-data selected genes had median 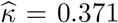. In the independent estimation halves, median 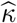 across the rank-based top-*K* sets ranged from 0.147 to 0.420; for the top-100 and top-250 analyses emphasized in Table 6, the range was 0.147–0.311. Because the selected gene sets differ between halves, this is not a paired estimate of selection bias, but the downward shift in most settings is consistent with winner’s-curse inflation in the same-sample audit. Strict half-sample discovery was also unstable: the two screening halves selected 0 and 14 genes at FDR 0.05, and the top-*K* lists had only 1%–2% Jaccard overlap. We therefore treat the full-data audit as descriptive and the top-100/top-250 analyses as independent-screening sensitivities rather than replicated discovery sets.

Per-gene 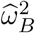 values were especially noisy in the independent estimation halves. Across the four top-*K* cutoffs, 16%–30% of genes in screen-1→estimate-2 and 4%–14% in screen-2→estimate-1 had *κ* ≤ 0; 28%–48% and 4%–25%, respectively, had 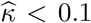. Correspondingly, 53%–60% and 32%– 46% of the gene-mean ratios fell outside [0, 1]. We therefore do not interpret medians or threshold counts of these cross-screen per-gene ratios as the headline result.

The pooled ratio-of-sums is much more stable. For screen-2→estimate-1, pooled gene-mean loss was 0.827 (0.762–0.886) for the top 100 genes and 0.836 (0.795–0.875) for the top 250, against weighted random benchmarks of 0.913 and 0.914. In the weaker screen-1→estimate-2 direction, pooled losses were 0.907 (0.842–0.969) and 0.932 (0.894–0.969), against benchmarks of 0.932 and 0.927. Thus one-dimensional gene means discard roughly 80%–93% of exposure-relevant information across the full and independent-screen analyses; they retain about twice the random-score information in the full data and the stronger cross-screen direction, but approach the random benchmark in the weaker direction. The two-component promoter-plus-non-promoter summary consistently improved the pooled loss: 0.625 in the full data and 0.634–0.714 across the four top-100/top-250 cross-screen analyses.

### 7.3 Independent-screening sensitivity

The independent-screening analysis therefore supports a collection-level adequacy statement, not stable identification of particular smoking-responsive genes and not precise gene-by-gene loss estimates. The final reproducibility script computes sign consistency exclusively in the screening half before evaluating adequacy in the opposite half, thereby avoiding conditioning on signs estimated from the same data used for 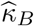 . That corrected sign-stratified analysis was heterogeneous across the two screening directions, so we treat it as exploratory and report it only in the reproducibility supplement rather than using it to explain the pooled result.

### 7.4 Outcome-specific expression illustration

We therefore studied expression of a gene as the outcome and the CpGs annotated to that gene as mediators, which keeps *p < n* so that IE_*M*_ is estimable by ordinary least squares. Of 417 adenocarcinomas with both methylation and RNA-seq (62 never-smokers), we screened eight candidate smoking-responsive genes by their adjusted total effect and analysed the three with the largest |*t*|: *AHRR* (83 CpGs, *t* = 2.93), *UCHL1* (10 CpGs, *t* = 2.48), and *NQO1* (9 CpGs, *t* = 2.14). *AHRR* is among the most consistently reported smoking-associated methylation loci (Zeilinger et al., 2013;Joehanes et al., 2016). Gene-level methylation was predictively associated with smoking after co-variate adjustment. The pre-specified primary aggregation was the gene mean; promoter means, complementary non-promoter means, a combined promoter-plus-non-promoter aggregation, a sign score, and one and three principal components were reported as secondary. Intervals are percentile intervals from a bootstrap stratified by smoking (2,000 resamples for *AHRR* and *UCHL1* ; 1,000 for *NQO1*), with sign weights and principal components re-estimated in each resample where indicated. Adequacy profiles are plug-in estimates on standardized CpGs.

Methylation and expression are measured concurrently, so the ordering of mediator and outcome is an assumption. Genes were selected on total effect from a short candidate list, and results should be read as an illustration of the audit rather than as new biology.

### 7.5 Outcome-specific results

For *AHRR*, the total effect was 0.63 (95% interval 0.27 to 0.99) and the CpG-level indirect effect 0.40 (−0.01 to 0.78). The gene-mean analysis gave −0.04 (−0.11 to 0.02), an interval that excludes any indirect effect larger than 0.02, and the gap was 0.44 (0.03 to 0.83) (Table 7, Figure 3). The non-promoter mean and the first principal component behaved identically. The gap for the gene mean remained between 0.44 and 0.45, with lower limits between 0.03 and 0.05, when stage, purity, or both were removed from the adjustment set. The CpG-level indirect effect itself was borderline under three of the four specifications; the robust finding is that the aggregate targets a different quantity, not that mediation is established. The promoter mean’s gap interval included zero. We call the complement “non-promoter” rather than “gene body” because the complementary annotation categories are not identical to the gene-body category. The canonical smoking-responsive *AHRR* CpG cg05575921 is annotated to the *AHRR* gene body in the Illumina 450K annotation and is a well-established smoking marker (Philibert et al., 2013; Joehanes et al., 2016; Maas et al., 2019). The sign score’s point estimate (0.39) was close to the CpG-level effect, but once its direction weights were re-estimated in each resample its interval widened to −0.20 to 0.77, illustrating the cost of estimating aggregation weights from the same exposure data.

**Table 7.** Application: indirect effects (log_2_ expression per smoking contrast) through CpG sets and their aggregates, main covariate specification. 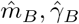 plug-in adequacy profile. Scores marked ^*†*^ were re-estimated in each bootstrap resample.

| Gene | Mediator | $k$ | IE<br>(95% interval) | $D_B$<br>(95% interval) | $\hat{m}_B$ | $\hat{\gamma}_B$ |
| --- | --- | --- | --- | --- | --- | --- |
| *AHRR* | All 83 CpGs | 83 | 0.40 (−0.01, 0.78) |  |  |  |
|  | Gene mean | 1 | −0.04 (−0.11, 0.02) | 0.44 (0.03, 0.83) | 0.84 | 0.39 |
|  | Promoter mean | 1 | 0.05 (−0.01, 0.12) | 0.35 (−0.04, 0.74) | 0.91 | 0.22 |
|  | Non-promoter mean | 1 | −0.04 (−0.12, 0.02) | 0.44 (0.03, 0.83) | 0.85 | 0.38 |
| | PC1$^\dagger$ | 1 | −0.04 (−0.11, 0.02) | 0.44 (0.03, 0.83) | 0.88 | 0.33 |
| | PC1–3$^\dagger$ | 3 | 0.10 (−0.05, 0.27) | 0.30 (−0.09, 0.68) | 0.73 | 0.46 |
| | Sign score$^\dagger$ | 1 | 0.39 (−0.20, 0.77) | 0.01 (−0.39, 0.58) | 0.48 | 0.74 |
| *UCHL1* | All 10 CpGs | 10 | 0.68 (0.21, 1.15) |  |  |  |
|  | Gene mean | 1 | 0.51 (0.08, 0.96) | 0.17 (−0.10, 0.43) | 0.25 | 0.47 |
|  | Promoter mean | 1 | 0.39 (−0.03, 0.81) | 0.29 (−0.00, 0.57) | 0.45 | 0.39 |
|  | Non-promoter mean | 1 | 0.57 (0.17, 0.99) | 0.11 (−0.18, 0.41) | 0.11 | 0.46 |
| | PC1$^\dagger$ | 1 | 0.50 (0.05, 0.95) | 0.18 (−0.07, 0.43) | 0.30 | 0.46 |
| | PC1–3$^\dagger$ | 3 | 0.46 (0.02, 0.91) | 0.22 (0.01, 0.42) | 0.13 | 0.34 |
| *NQO1* | All 9 CpGs | 9 | 0.08 (−0.17, 0.32) |  |  |  |
|  | Gene mean | 1 | −0.01 (−0.18, 0.17) | 0.08 (−0.08, 0.28) | 0.98 | 0.09 |
|  | Promoter mean | 1 | −0.01 (−0.15, 0.15) | 0.08 (−0.10, 0.29) | 0.97 | 0.13 |
| | PC1$^\dagger$ | 1 | −0.02 (−0.17, 0.16) | 0.09 (−0.08, 0.30) | 0.94 | 0.24 |

**Figure 3.**
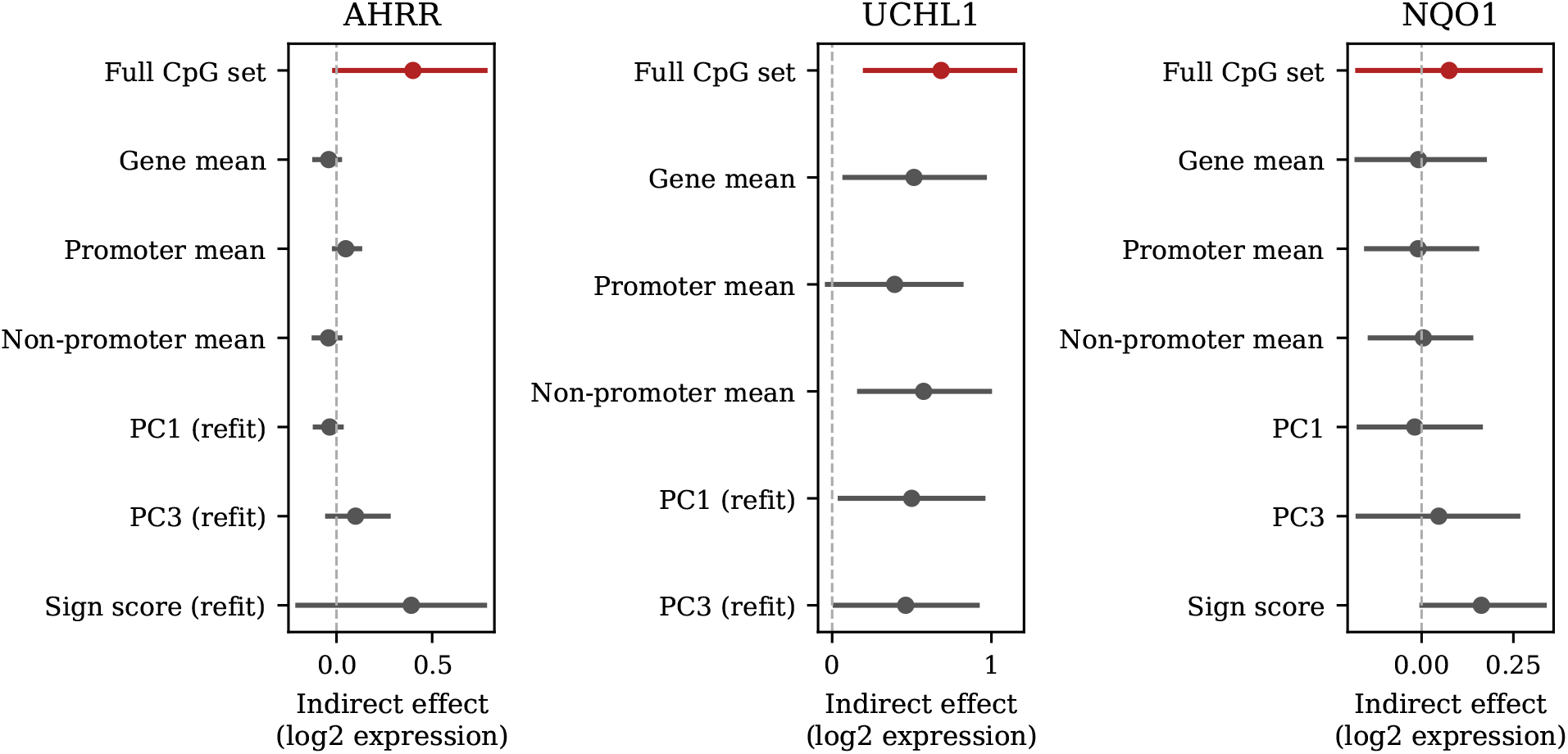
Indirect effects of smoking on gene expression through all annotated CpGs and through common aggregates, with 95% stratified bootstrap intervals. *Alt text:* Interval plot for AHRR, UCHL1, and NQO1 comparing the CpG-level indirect effect with gene means, promoter/non-promoter means, principal components, and sign scores. AHRR shows the clearest separation between the full CpG effect and several one-score aggregates; UCHL1 aggregates generally attenuate the full effect; NQO1 intervals include zero.

For *UCHL1*, the CpG-level indirect effect was 0.68 (0.21 to 1.15) and was significant under all specifications (lower limits 0.15 to 0.26). Every aggregation also detected mediation but understated it: the promoter mean by 43% (gap 0.29; −0.00 to 0.57 in the main specification, excluding zero in two of four specifications). For *NQO1*, neither the CpG-level analysis (0.08; −0.17 to 0.32) nor any aggregation indicated mediation. The reported *NQO1* PC1 interval is from the seed-11 robustness run in which the principal-component weights are re-estimated within each bootstrap resample; it is therefore marked as a learned-score sensitivity.

For *AHRR* and *UCHL1*, all 13 observed relative gaps lay within the plug-in half-range 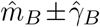 for example, the *AHRR* promoter mean had observed relative gap 0.88 against 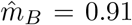, and the *UCHL1* promoter mean 0.43 against 0.45. Because aggregations within a gene share a single realization of the outcome effects, and because plug-in 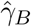 is biased upward, this agreement is consistency rather than validation. For *NQO1*, the indirect effect is too close to zero for relative gaps to be meaningful.

## 8 Discussion

Aggregating mediators changes the mediated estimand. The change has a simple geometry: *B* retains one projection of the exposure and outcome directions, the omitted inner product becomes *D*_*B*_, and that same quantity reappears in the direct effect. The total effect is therefore preserved even when the mediation decomposition is badly distorted. This explains why conventional internal checks—a stable total effect, a well-fit aggregate regression, or a narrow interval around IE_*Z*_— cannot establish that a pathway score represents mediation through its components.

The extended framework separates universal, benchmark, and outcome-specific adequacy. Universal adequacy is characterized exactly by Σ^*−*1^*α* ∈ col(*B*) and summarized continuously by *ω*_*B*_. Benchmark adequacy is described by (*m*_*B*_, *γ*_*B*_) and the Cauchy tail probabilities, which require no outcome observations but do require a reference distribution over outcome directions. Outcome-specific adequacy is the realized *D*_*B*_, for which Theorem 2 supplies formal paired inference. These three layers answer different questions and should not be collapsed into a single “valid/invalid” label.

The minimax result sharpens the adequacy–overlap trade-off. The identity 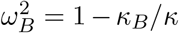 says that the same exposure information discarded to improve overlap is exactly the information whose removal creates worst-case vulnerability to mediated-effect distortion. There is therefore no free dimension reduction that simultaneously preserves every possible mediated effect and makes the exposure substantially less predictable from the mediator set. This conclusion is distribution-free within the linear projection model and does not depend on the spherical benchmark used for the Cauchy profile.

The profile contributes a different form of information. Its spherical generalization shows that normality of *β* was stronger than necessary; only random direction, not radial magnitude, drives the Cauchy ratio. Sparse outcome effects in the semi-synthetic study are far from spherical, yet the location and half-interquartile range remained close to the theoretical profile. That robustness should not be overinterpreted. Biological outcome effects may be concentrated in pathways that align systematically with either retained or omitted mediator directions. When substantive information about that anisotropy exists, the benchmark distribution should encode it rather than default to spherical symmetry.

Formal gap inference addresses an important limitation of purely pre-outcome diagnostics. Because *D*_*B*_ = DE_*Z*_ − DE_*M*_, no new causal parameter has to be estimated in moderate dimensions; the key is to use the paired covariance. The finite-sample study shows that treating the two direct-effect estimates as independent can be dramatically conservative. The test also prevents a common interpretive error: a gene or pathway score can have an indirect effect close to zero with a tight interval while the component-level indirect effect is materially different, as in the *AHRR* illustration. In that setting, the relevant inferential object is not only whether IE_*Z*_ differs from zero but whether IE_*Z*_ differs from IE_*M*_ .

Several practical conclusions follow from the TCGA analyses. Most importantly, the genome-wide audit makes the aggregation problem a collection-level finding rather than a three-gene anecdote, but the appropriate headline is the pooled ratio of information sums rather than a median of unstable gene-specific ratios. In the full-data audit the gene mean discarded 81.1% of pooled exposure-relevant information (95% descriptive gene-bootstrap interval 79.7%–82.3%), compared with a 91.3% dimension-only random benchmark. Independent screening sharply reduced discovery stability and estimated exposure information, consistent with a winner’s-curse component in the same-sample audit; nevertheless, top-100/top-250 cross-screen pooled losses remained 82.7%– 93.2%. The stronger cross-screen direction retained roughly twice the information of a dimension-matched random score, whereas the weaker direction approached the random benchmark. Keeping promoter and non-promoter components separate reduced pooled loss to 62.5% in the full data and 63.4%–71.4% across the independent sensitivities. Because these audits were completed without introducing expression, the conclusions cannot be attributed to selecting genes whose downstream outcomes happened to favor the full-mediator model.

The audit also clarifies what should *not* be concluded. Cross-screen gene identities are highly unstable, and many estimation-half 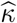 values are close to or below zero; individual 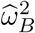ratios are therefore too noisy to support statements such as “most specific genes lose more than half.” The robust statement concerns pooled information across prespecified collections. Exposure weighting is not automatically protective; the direction *B* = *α* can have *m*_*B*_ = 0 yet a large *γ*_*B*_ because correlation rotates the exact exposure-predictive direction to Σ^*−*1^*α*. Retaining more principal components generally reduces distortion, but the number required can be large and the four-fold profile estimator becomes unreliable when *k* is a substantial fraction of *n*. Data-learned weights also require an additional layer of uncertainty. The *AHRR* sign score illustrates this sharply: its point estimate nearly reproduces the full CpG-level effect, while refitting the sign directions inside the bootstrap produces a much wider interval. Finally, annotation-based summaries need precise labels; “non-promoter” is not synonymous with “gene body.”

The paper has limitations. The structural theory is linear and homogeneous. With exposure– mediator interactions, nonlinear outcome regressions, or non-collapsible links, the aggregation gap can depend on the exposure contrast and the projection identity will not carry over unchanged. Stochastic-intervention mediation effects (Díaz and Hejazi, 2020) offer one route to an extension. The fixed-dimensional gap theorem does not solve inference when *p* is comparable with or greater than *n*; Section 4.2 deliberately treats DML as a route for future theory rather than as an established result of this paper. The profile estimator itself has a practical range: in the present design, *k* up to roughly *n/*20 was reliable, whereas larger subspaces produced unstable 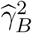 . The genome-wide audit is outcome-free but still comes from one tumour tissue, one array platform, and one observational exposure contrast; transport to blood, EPIC methylation, other omics layers, or other exposures remains to be established. The full-sample gene screen and adequacy estimates use the same participants, and the independent half-sample analysis shows both that smoking-responsive gene discovery is highly unstable at roughly *n* = 210 per screening half and that same-sample screening can inflate estimated exposure information. The cross-screen top-*K* analysis therefore supports a collection-level sensitivity conclusion, not stable identification of particular smoking-responsive genes. Untruncated gene-specific 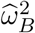values are especially unstable when 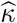 is close to zero; this is why the pooled ratio-of-sums, together with the random-subspace benchmark, is primary and individual-gene ratios are secondary. The gene-expression illustration uses concurrently measured tumour methylation and expression, so temporal mediation is assumed rather than demonstrated, and the three detailed genes were selected from an eight-gene candidate list using total-effect statistics. Those outcome-specific results remain illustrations of estimand distortion rather than new evidence establishing a biological mediation mechanism.

A stronger external validation would freeze *B* prospectively in a cohort where exposure precedes methylation and methylation precedes expression or a later phenotype. Epigenetic clocks are especially attractive because the weights are published in advance (Horvath, 2013; Lu et al., 2019). Replicating the outcome-free audit in a second tissue or array would test whether the high loss of one-number gene summaries is a feature of smoking-responsive methylation more broadly rather than of TCGA LUAD specifically.

The main recommendation is therefore modest: before a composite mediator is given the causal interpretation of its components, audit the aggregation. Report what fraction of the exposure-relevant mediator direction it retains, how large its relative gap can plausibly be under a transparent benchmark, and—when the full effect is estimable—whether the observed gap is statistically distinguishable from zero. A low-dimensional score can still be scientifically useful even when it fails these checks, but its indirect effect should then be described as mediation through the score itself rather than through the full collection it summarizes.

## Software and data availability

TCGA data are publicly available through UCSC Xena. Reproducibility code and frozen reference outputs for the simulations, genome-wide audit, independent screen/estimate analyses, and empirical application are available at GitHub and are archived as version 1.0.2 on Zenodo (doi:10.5281/zenodo.23106680). The public archive reproduces the analyses from the processed TCGA LUAD stage-1 object onward. The processed stage-1 object is not redistributed, and the exact historical raw-data preprocessing script that created it is not included in the recoverable analysis archive.

## Appendix A Proofs

### Proof of Proposition 1

Let 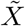 denote the residual of *X* after linear projection on *W* . Then 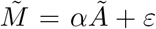 and, since *ε* is uncorrelated with 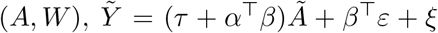. Decompose 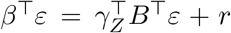SSwith *γ*_*Z*_ = (*B*^*⊤*^Σ*B*)^*−*1^*B*^*⊤*^Σ*β*, so that *r* is uncorrelated with *B*^*⊤*^*ε* and with (*A, W*). Substituting 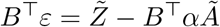gives

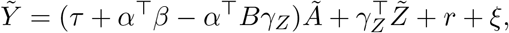

where *r* + *ξ* is uncorrelated with (*Ã*, 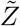). Hence DE_*Z*_ = *τ* + *α*^*⊤*^*β* − *α*^*⊤*^*P*_*B*_*β*, and IE_*Z*_ = TE − DE_*Z*_ = *α*^*⊤*^*P*_*B*_*β*. The remaining identities follow. □

### Proof of Proposition 2

(i) *D*_*B*_ = 0 for all *β* iff (*I* − *P*_*B*_)^*⊤*^*α* = 0, i.e. *α* = Σ*B*(*B*^*⊤*^Σ*B*)^*−*1^*B*^*⊤*^*α* ∈ col(Σ*B*), i.e. Σ^*−*1^*α* ∈ col(*B*). Conversely, if *α* = Σ*Bc* then 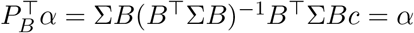. (ii) *D*_*B*_ = 0 for all *α* iff *P*_*B*_*β* = *β*; since range(*P*_*B*_) = col(*B*) and *P*_*B*_*B* = *B*, this holds iff *β* ∈ col(*B*). (iii) *α*^*⊤*^*P*_*B*_*β* = *a*^*⊤*^Π_*B*_*b*, and since Π_*B*_ is a symmetric idempotent, *a*^*⊤*^(*I* −Π_*B*_)*b* = ⟨(*I* −Π_*B*_)*a*, (*I* −Π_*B*_)*b*⟩.

□

### Proof of Proposition 3

Conditional on *W* = *w*, Bayes’ rule gives

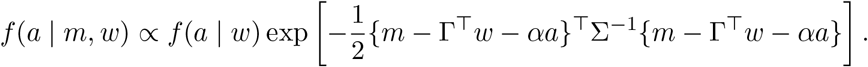

Terms involving both *a* and *m* enter only through *a α*^*⊤*^Σ^*−*1^(*m* − Γ^*⊤*^*w*). Hence the conditional distribution of *A* given (*M, W*) depends on *M* only through *T* (*M, W*), proving sufficiency by factorization. Under a non-degenerate one-index inverse model, the associated central subspace is therefore span(Σ^*−*1^*α*). Proposition 2(i) then gives the final equivalence with universal mediation preservation. □

### Proof of Proposition 4

With the notation above, 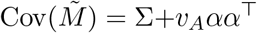 and 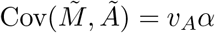. By the Sherman–Morrison formula the projection coefficient is (Σ+ *v*_*A*_*αα*^*⊤*^)^*−*1^*v*_*A*_*α* = *v*_*A*_Σ^*−*1^*α/*(1 +*v*_*A*_*κ*), and the residual variance is *v*_*A*_ − *v*_*A*_*α*^*⊤*^(·) = *v*_*A*_*/*(1 + *v*_*A*_*κ*). Replacing 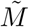 by 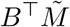, with covariance *B*^*⊤*^Σ*B* + *v*_*A*_*B*^*⊤*^*αα*^*⊤*^*B*, gives *v*_*A*_*/*(1 + *v*_*A*_*κ*_*B*_). Finally *κ*_*B*_ = *a*^*⊤*^Π_*B*_*a* ≤ *a*^*⊤*^*a* = *κ*, with equality iff *a* ∈ col(Σ^1*/*2^*B*), i.e. Σ^*−*1^*α* ∈ col(*B*). □

### Proof of Corollary 2

(i) If *α* = *Bc* then *P*_*B*_*α* = *α*, so *α*^*⊤*^*u* = ∥*α*∥^2^ − *α*^*⊤*^*P*_*B*_*α* = 0. (ii) *γ*_*B*_ = 0 iff *u* = *cα* for some *c*, in which case *m*_*B*_ = *c*; both vanish iff *u* = 0, which is Proposition 2(i). (iii) If Σ*B* = *B*Λ;_*k*_ with *B*^*⊤*^*B* = *I*_*k*_, then 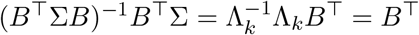 so *P*_*B*_ = *BB*^*⊤*^ is symmetric, *u* = (*I* − *BB*^*⊤*^)*α, α*^*⊤*^*u* = ∥*u*∥^2^, and *m*_*B*_ = ∥*u*∥^2^*/*∥*α*∥^2^. Then 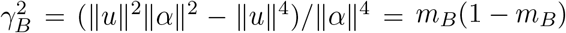. Rescaling the eigenvalues leaves *B* and hence *P*_*B*_ unchanged. □

### Proof of Proposition 6

Conditional on the two design matrices, Gaussian multivariate regression gives independence between each OLS coefficient vector and its residual sum-of-squares matrix, and the two splits are independent. The pooled residual sum of squares satisfies

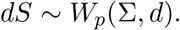

Therefore *B*^*⊤*^(*dS*)*B* ∼ *W*_*k*_(*B*^*⊤*^Σ*B, d*). For *d > k* + 1, the inverse-Wishart moment gives

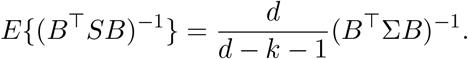

Also 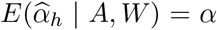, and the coefficient vectors are independent of *S* and of each other across splits. Taking expectations in (6) thus yields *α*^*⊤*^*B*(*B*^*⊤*^Σ*B*)^*−*1^*B*^*⊤*^*α* = *κ*_*B*_. Setting *B* = *I*_*p*_ gives the second result with the factor (*d* − *p* − 1)*/d*. □

### Proof of Proposition 7

For *B*^*\**^ = *BR* with invertible *R*,

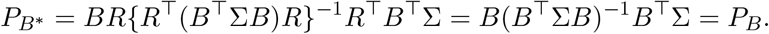

All quantities listed in part (i) are functions of this projector or its corresponding whitened subspace. For part (ii), nested column spaces imply nested orthogonal projection subspaces for 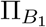 and 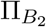 . Orthogonal projection onto the larger subspace cannot increase the residual norm, so 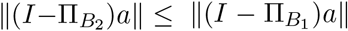. Divide by ∥*a*∥ to obtain the ordering of *ω*_*B*_. Because *κ*_*B*_ = ∥Π_*B*_*a*∥^2^, the reverse ordering holds for *κ*_*B*_. □

### Proof of Proposition 5

Because *a* = Σ^*−*1*/*2^*α* and Π_*B*_ is an orthogonal projector,

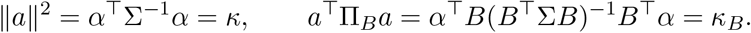

Pythagoras gives ∥(*I* − Π_*B*_)*a*∥^2^ = *κ* − *κ*_*B*_, proving (3). Further,

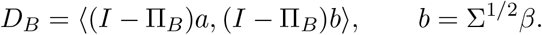

If *β*^*⊤*^Σ*β* = ∥*b*∥^2^ ≤ 1, Cauchy–Schwarz gives |*D*_*B*_| ≤ ∥(*I* − Π_*B*_)*a*∥. Equality is attained by taking *b* proportional to (*I* − Π_*B*_)*a* when this vector is nonzero. Dividing by 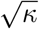 proves (4). Finally Proposition 4 implies 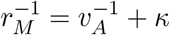 and 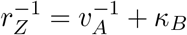 substitution into (3) gives (5). □

### Proof of Theorem 1

Write *β* = *RU* with *R >* 0 independent of *U* and *U* uniform on the unit sphere. Let *G* ∼ *N* (0, *I*_*p*_). Since 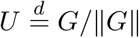

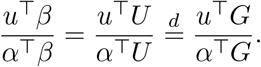

The pair (*α*^*⊤*^*G, u*^*⊤*^*G*) is centred bivariate normal with variances ∥*α*∥^2^, ∥*u*∥^2^ and covariance *α*^*⊤*^*u*. Decompose *u*^*⊤*^*G* = (*α*^*⊤*^*u/*∥*α*∥^2^)*α*^*⊤*^*G* + *E*, where *E* is normal, independent of *α*^*⊤*^*G*, with variance ∥*u*∥^2^ −(*α*^*⊤*^*u*)^2^*/*∥*α*∥^2^. The ratio of the independent centred normals is standard Cauchy after scaling, yielding the stated *m*_*B*_ and *γ*_*B*_. If the residual variance is zero the ratio is degenerate at *m*_*B*_. □

### Proof of Theorem 2

For either adjustment set *s*, the Frisch–Waugh–Lovell representation is

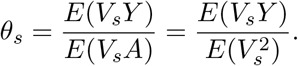

The ordinary least-squares coefficient is an *M* -estimator whose first-order expansion for the coefficient on *A* is

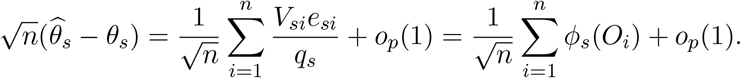

The two estimates are computed on the same observations, so subtracting the two expansions rather than combining marginal variances gives

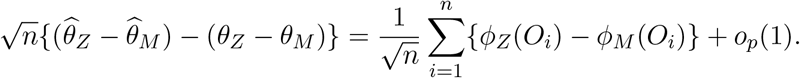

The central limit theorem gives the stated limit under the moment assumptions. Replacing population residuals and *q*_*s*_ by their least-squares sample analogues is consistent under fixed dimension, so the empirical variance of the estimated influence-function differences consistently estimates *V*_*D*_. Proposition 1 supplies *θ*_*Z*_ − *θ*_*M*_ = *D*_*B*_. □

## B Supplementary results

### Constructed outcome scenarios

Table 8 gives IE_*Z*_ in four constructed scenarios with 50 (or 100) nonzero outcome effects of size 0.02 on standardized CpGs: effects aligned in sign with the 50 largest exposure effects; alternating signs across the 100 largest (cancellation); effects on 50 hypermethylated CpGs only; and effects of size 0.10 on the 50 CpGs with the smallest exposure effects. In the last scenario the true indirect effect is 0.010, but PC1 reports −0.31 and the hyper/hypo means report 0.13: aggregates borrow exposure signal from correlated CpGs that have no outcome effect.

**Table 8.**
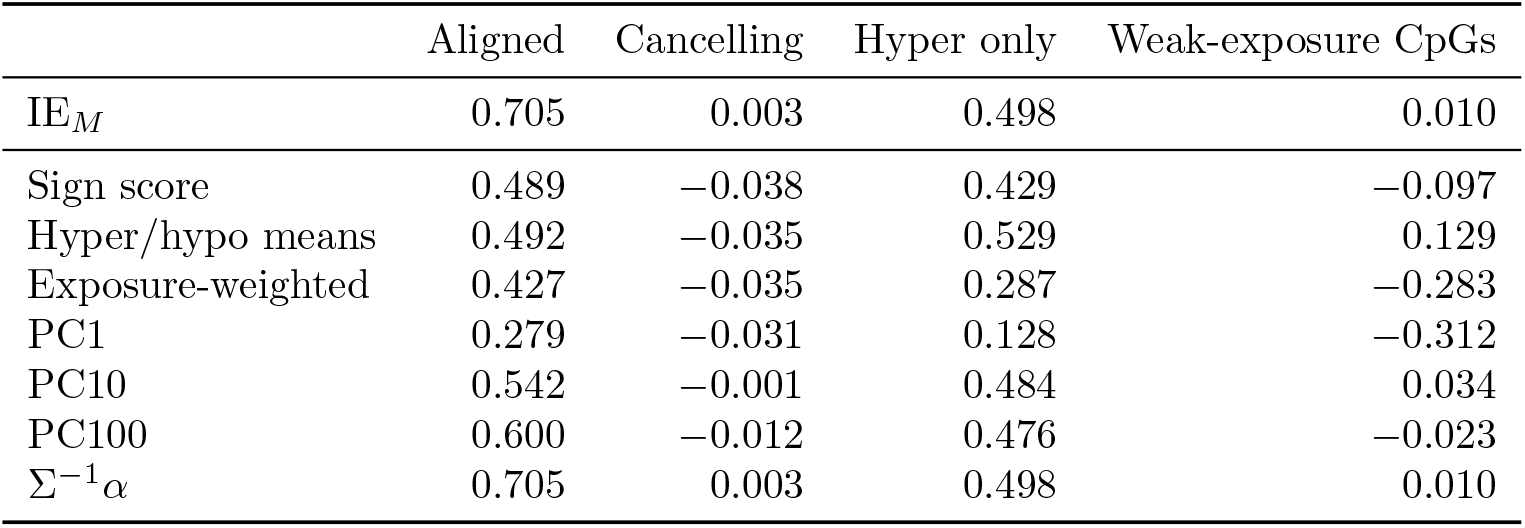
True IE_*Z*_ in constructed scenarios (population, *δ* = 0.2).

### Robustness of the application

Table 9 reports the gene-mean gap and the CpG-level indirect effect under four adjustment sets.

**Table 9.** Sensitivity of the application to the adjustment set (2,000 stratified bootstrap resamples).

| Gene | Adjustment set | $\text{IE}_M$ (95% interval) | Gene-mean $D_B$ (95% interval) |
| --- | --- | --- | --- |
| *AHRR* | Age, sex, stage, purity | 0.40 (-0.01, 0.78) | 0.44 (0.03, 0.83) |
|  | Age, sex, purity | 0.40 (-0.00, 0.80) | 0.44 (0.04, 0.83) |
|  | Age, sex, stage | 0.41 (-0.00, 0.80) | 0.45 (0.03, 0.84) |
|  | Age, sex | 0.41 (0.00, 0.82) | 0.45 (0.05, 0.85) |
| *UCHL1* | Age, sex, stage, purity | 0.68 (0.21, 1.15) | 0.17 (-0.10, 0.43) |
|  | Age, sex, purity | 0.63 (0.15, 1.10) | 0.14 (-0.13, 0.40) |
|  | Age, sex, stage | 0.72 (0.26, 1.18) | 0.21 (-0.04, 0.46) |
|  | Age, sex | 0.66 (0.18, 1.12) | 0.17 (-0.08, 0.41) |

## C Supplementary implementation details

### C.1 Gap-inference simulation construction

For reproducibility, the gap-inference study uses a deterministic population setup for each *p*. The mediator residual covariance has entries Σ_*jk*_ = 0.35^|*j−k*|^, and *B* = *p*^*−*1*/*2^**1**. Exposure effects decay smoothly with index and alternate modestly in sign every third component. Under the null, *β* is proportional to *B* and scaled to give IE_*M*_ = 0.35, which guarantees *D*_*B*_ = 0 by Proposition 2(ii). Under the alternative, {0.20*/*(*u*^*⊤*^*u*)}*u* is added to *β*, with *u* = (*I* − *P*_*B*_)^*⊤*^*α*, giving *D*_*B*_ = 0.20 algebraically. The base-R script fixes random seed 16092026, uses 800 replicates per cell, and writes the exact table source to `gap_inference_results_R.csv`.

### C.2 A reporting template for aggregation audits

A reproducible report should identify (1) the scientific unit represented by each column of *M* and its scale; (2) whether *B* was prespecified, imported from an external source, or learned in the analysis sample; (3) the rank *k* and any normalization of *B*; (4) the overlap quantities *r*_*M*_ and *r*_*Z*_ when estimable; (5) *ω*_*B*_ or an explanation of why it cannot be stably estimated; (6) (*m*_*B*_, *γ*_*B*_) and at least one threshold probability; (7) IE_*M*_, IE_*Z*_, and the paired interval for *D*_*B*_ when the full mediator model is estimable; and (8) the exact resampling rule used for learned scores. These items distinguish score construction, structural adequacy, and sampling uncertainty.

For principal components, the covariance matrix from which the loadings were learned must be stated. Corollary 2(iii) requires eigenvectors of the residual covariance Σ appearing in the mediator model. PCs from the raw CpG matrix, from an exposure-stratified matrix, or from a separately standardized feature matrix need not obey 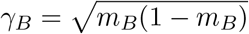. For annotation-based means, the annotation rule should be reported explicitly; complements should not be relabeled as a biological region unless the annotation supports that label.

## Notes

https://xena.ucsc.edu/

https://github.com/subirhait/pathway-score-mediation

https://doi.org/10.5281/zenodo.23106680

### Competing Interest Statement

The authors have declared no competing interest.

